# Standing Variations Modeling Captures Inter-Individual Heterogeneity in a Deterministic Model of Prostate Cancer Response to Combination Therapy

**DOI:** 10.1101/2021.02.04.429807

**Authors:** Harsh Vardhan Jain, Inmaculada C Sorribes, Samuel K Handelman, Johnna Barnaby, Trachette L Jackson

## Abstract

Sipuleucel-T (Provenge) is the first live cell vaccine approved for advanced, hormonally refractive prostate cancer. However, survival benefit is modest and the optimal combination or schedule of sipuleucel-T with androgen depletion remains unknown. We employ a nonlinear dynamical systems approach to modeling the response of hormonally refractive prostate cancer to sipuleucel-T. Our mechanistic model incorporates the immune response to the cancer elicited by vaccination, and the effect of androgen depletion therapy. Because only a fraction of patients benefit from sipuleucel-T treatment, inter-individual heterogeneity is clearly crucial. Therefore, we introduce our novel approach, Standing Variations Modeling, which exploits inestimability of model parameters to capture heterogeneity in a deterministic model. We use data from mouse xenograft experiments to infer distributions on parameters critical to tumor growth and to the resultant immune response. Sampling model parameters from these distributions allows us to represent heterogeneity, both at the level of the tumor cells and the individual (mouse) being treated. Our model simulations explain the limited success of sipuleucel-T observed in practice, and predict an optimal combination regime that maximizes predicted efficacy. This approach will generalize to a range of emerging cancer immunotherapies.

## 1. Introduction

Prostate cancer (PCa) is the most common malignancy affecting men in the US, and a leading cause of cancer related deaths [1]. PCa also contributes to racial health disparities, as lethality in men of recent African ancestry is roughly twice as high as in men of European ancestry [2]. Typically, the first-line treatment for patients with biochemically failing and metastatic PCa involves blocking the bioavailability of androgens to cancer cells by the constant or periodic application of a combination of chemical castration agents. This is known as androgen deprivation therapy (ADT). However, a majority of men treated with ADT progress to a castration resistant state, necessitating alternative interventions [3]. Therapeutic advances made over the last decade have conferred varying degrees of survival benefit to patients with metastatic castration-resistant prostate cancer (mCRPC) [4].

Sipuleucel-T (Provenge) is an autologous cellular immunotherapy approved by the FDA in 2010 for the treatment of patients with mCRPC [4]. Sipuleucel-T is a PCa live cell vaccine consisting of a patient’s own peripheral blood mononuclear cells (PBMC). PBMC are harvested via leukapheresis, and cultured in the presence of a recombinant protein, a chimera of prostatic acid phosphatase (PAP) and granulocyte-macrophage colony-stimulating factor (GM-CSF). This chimera activates the mononuclear cells, transforming them into antigen presenting cells (APCs) which are re-injected into the patient 3 days later [4,5].

However, sipuleucel-T has not been widely adopted [6]. This is due, in part, to the limited success it has exhibited clinically. The first phase III clinical trial of sipuleucel-T reported only a modest median overall survival benefit of 4.1 months (25.8 versus 21.7 months over the placebo group), and no significant difference in time to progression (3.7 versus 3.6 months over the placebo group) [7]. Furthermore, the vaccine is given in three doses typically administered every two weeks [4]. But, as the FDA notes [8], an optimal schedule has not been established, with intervals between doses ranging from 1 to 15 weeks in controlled clinical trials.

Our aim is to uncover processes governing the response of cancer cells and individual patients (here, mice) to immunotherapy. From these processes, mathematical models will recapitulate clinical results and predict dosing schedules and therapy combinations with better outcomes. To this end, we develop a nonlinear dynamical systems model of prostate cancer response to sipuleucel-T, alone or in combination with ADT, which incorporates a wide range of biological interactions and processes. We calibrate our model with experimental data on the treatment of PCa xenografts in mice, reported in [9]. Consequently, (virtual) mice are used here as a surrogate for human patients.

We next introduce our novel modeling paradigm, *Standing Variations Modeling*, which exploits uncertainty in parameter values to derive information on the heterogeneity that is a characteristic feature of cancers. This heterogeneity is key to understanding the results of the sipuleucel-T clinical trial, and informs alternative strategies for maximizing the therapeutic potential of this vaccine. In our approach, we first infer probability distributions from which model parameters most likely arise. This differs from traditional modeling approaches that only use static or mean expression levels and cellular responses, ignoring variation across cell populations or patients. We then sample randomly from these posterior distributions on model parameters, resulting in parameter vectors, each of which can be thought of as a distinct individual within a (virtual) population. The simulated cohort of heterogeneous individuals is then used for treatment optimization and conducting *in silico* clinical trials. Our Standing Variations Model is inspired by approaches in evolutionary biology [10,11], where inter-individual (inter-patient, or inter-rodent) differences arise from existing diversity in the population; in our paradigm, this diversity may or may not have a genetic basis.

We build on previously proposed mathematical models of PCa response to ADT [12–21], as well as models of tumor-immune interactions (we refer the reader to [22,23] for recent reviews). Recent studies have also reported models of PCa response to vaccination alone [24,25] or in combination with ADT [26]. Our approach differs from these in three major respects. Our model of PCa and immune system interactions builds on the tumorimmune interaction models of Radunskaya et al. [27] and Robertson-Tessi et al. [28], and on Rutter and Kuang’s model of PCa treatment with ADT and immunotherapy [26] by incorporating a higher level of biological detail, tailored to the specifics of PCa. We will show that these additional relationships drive emergent behaviors important to the outcome of PCa and ADT combination therapy. Moreover, terms in our model are mechanistically derived from bio-molecular first principles, whenever possible. Finally, in addition to utilizing experimental data for parameter estimation, our Standing Variations approach reformulates this inverse problem by inferring the distributions from which model parameters arise. Thus, we are able to capture variability within, and across individuals with a deterministic model.

## 2. Materials and Methods

### 2.1. Model schematic

Our model of PCa growth and response to ADT and immunotherapy is cast as a system of coupled non-linear ordinary differential equations (ODEs) tracking the temporal evolution of key cellular and chemical species. A model schematic is shown in Figure 1.

**Figure 1.**
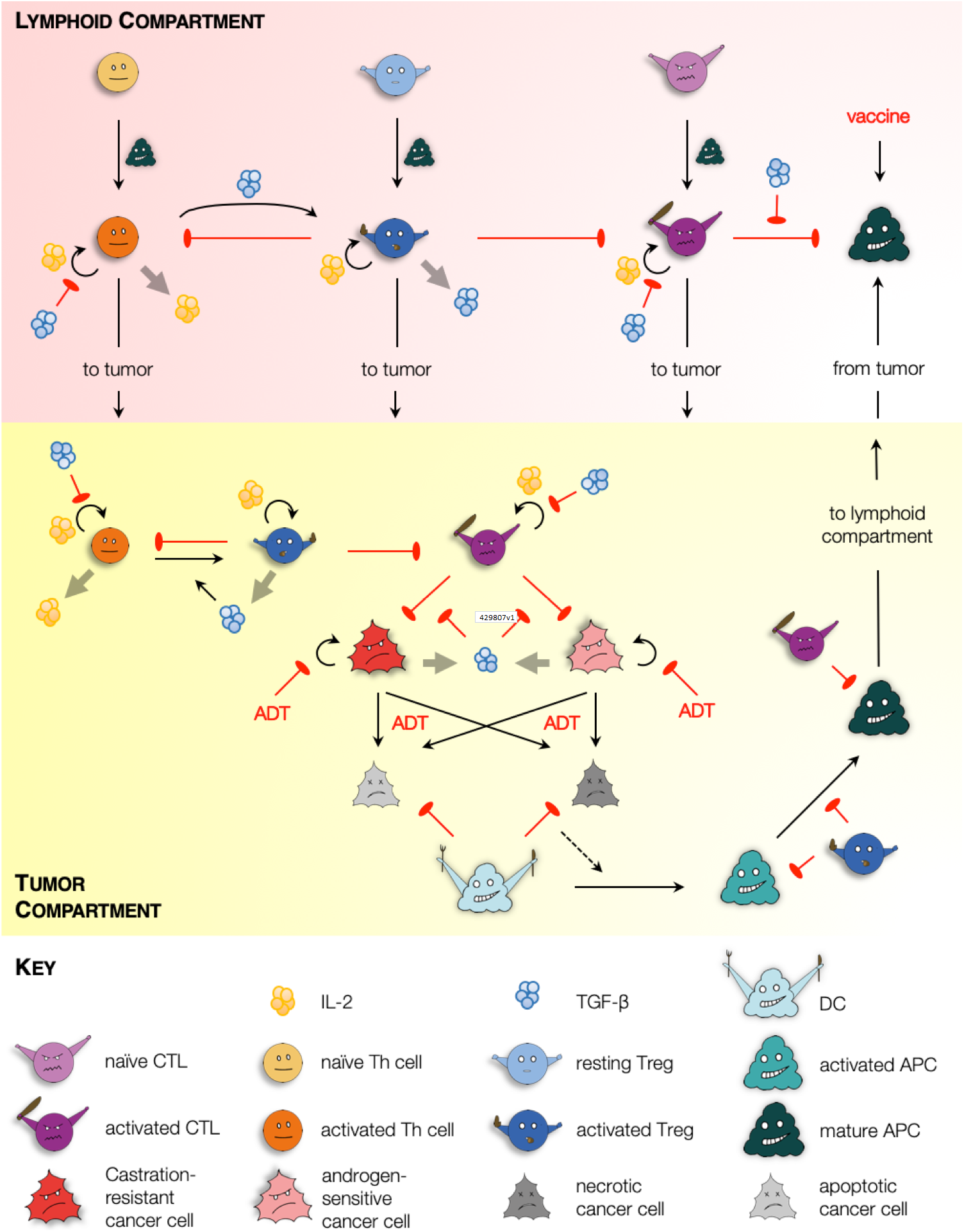
Model schematic. Solid black arrows indicate transformation of one species to another, or transport of species. Semi-circular black arrows indicate proliferation. Red arrows with flat heads indicate inhibition or cell kill. Light grey arrows indicate expression of chemokines.

In our formulation, two physiological compartments are considered where model species reside: the tumor; and lymphatic organs such as tumor draining lymph nodes or the spleen. Within the tumor, our model captures androgen-sensitive and castration-resistant cancer cell proliferation, and two forms of cell death, namely, apoptosis and necrosis, which occur in response to environmental stresses or applied treatment [29]. Dead cells are cleared from the tumor site by phagocytes such as macrophages (not included explicitly in our model) and, to a lesser extent, dendritic cells (DCs) [30]. Specifically, dying cells in our model release ‘find-me’ signaling molecules such as LysoPC and S1P, which trigger an innate immune response resulting in the recruitment of immature DCs to the tumor site [31]. We make the important distinction between cell death via apoptosis and necrosis because immature DCs can only transform into activated antigen presenting cells (APCs) when they phagocytose necrotic tumor cells [32]. Our model also captures APC maturation within the tumor along with their transformation to a phenotype marked by the up-regulation of co-stimulatory molecules such as CD80/CD86, and the expression of various cytokines necessary for the activation of effector T cells [33].

Mature APCs subsequently migrate to lymphoid organs such as tumor draining lymph nodes or the spleen where they activate resident naïve T cells into CD4^+^ helper T (Th) cells and CD8^+^ cytotoxic T lymphocytes (CTLs) [33,34]. Activated Th cells, in turn, secrete cytokines such as IL-2 which induces proliferation of all activated T cell populations [35]. Post-activation, our model assumes that all activated T cell populations migrate to the tumor site [34]. To account for fact that the speed of CTL infiltration deceases as tumor volume increases [36], the maximum rate of CTL-induced apoptosis is scaled by the radius of the tumor. Within both, the tumor and the lymphoid compartments, CTLs induce cell death via direct contact in any cell type that expresses their cognate antigen, for instance tumor cells, and activated and mature APCs [37].

We also include regulatory T cells (Tregs) in our framework. Two primary types of Tregs have been identified *in vivo*, namely, naturally occurring or thymus-derived Tregs (tTregs) and peripheral Tregs (pTregs) [38]. tTregs develop in the thymus and may further be classified as resting and activated [39]. The activation of tTregs depends on T cell receptor (TCR) stimulation [38,39], for instance, by mature APCs [40]. On the other hand, pTregs are generated *de novo* from conventional CD4^+^ T cells [41], including at the site of the tumor [42]. Indeed, there is evidence that an increase in Treg numbers in cancer patients is a consequence of both, the recruitment of tTregs [42], and the generation of pTregs [41,42]. We account for these processes in our model as follows. For simplicity, we do not distinguish between tTregs and pTregs; rather, resting Tregs are assumed to localize at a constant rate to the lymphoid compartment, where they undergo activation after coming in contact with mature APCs. Additionally, activated Th cells may transform directly into activated Tregs as described below. Once activated, Tregs migrate to the tumor site [43].

Within both, the tumor and the lymphoid compartments, activated Tregs are assumed to exert their immunosuppressive functions in two distinct ways: (i) long range interactions, via the expression of immunosuppressive cytokines such as IL-10 and TGF-*β*; and (ii) short range interactions involving direct contact with target cells [38,44,45]. Here, we explicitly include TGF-*β* expression and function, although the model readily generalizes to including additional chemokines. TGF-*β* induces the conversion of activated CD4^+^ T cells into activated Treg cells [43,46,47]. This conversion may also occur within the tumor, under signaling from TGF-*β* secreted by cancer cells, macrophages and activated Tregs [45]. Since we do not model macrophages explicitly, we assume that the source term for macrophage-derived TGF-*β* is proportional to the numbers of dead cells. TGF-*β* also exerts an immunosuppressive effect by inhibiting: the production of IL-2 by activated Th cells; the proliferation of activated Th cells and CTLs; and CTL cytotoxicity [48]. Additionally, Tregs down-regulate the activation of the transcription factor NF-*κ*B in DCs, thereby inhibiting their maturation [49]. All of these processes are incorporated in our model. Finally, activated Tregs induce cell death in effector T cells or CTLs in a contact-dependent manner involving granzymes A and B [50].

The effects of ADT and vaccination with sipuleucel-T are incorporated as follows. Under ADT, androgen sensitive cancer cells cease to proliferate and undergo cell death primarily via apoptosis, although some cells may undergo necrosis or necroptosis [51]. Castration resistant cancer cells are assumed to be dependent on androgens to a far lesser extent; consequently, their proliferation rate may slow down under ADT, and limited cell death may be induced. Under vaccination, mature APCs are injected into circulation, from where they may undergo clearance or accumulate in highly vascularized organs such as the spleen [52]. Once in the spleen, the mature APCs are presumed to trigger an anti-tumor immune response as described above.

Further details on model derivation and the full system of ODEs representing the above processes are available in the Supplementary Information.

### 2.2. Parameter estimation

Where possible, parameter values were taken from the literature. Values for the remaining parameters were chosen so that model simulations matched available experimental data, in a least squares sense. The results of these fits are shown in Figure 2. In particular, parameters relating to tumor cell proliferation and death, dead cell clearance, immune cell infiltration into the tumor, and chemokine expression were fit to data from PCa xenograft experiments reported in [9]. Rates of tumor cell apoptosis and necrosis were estimated from data reported in [53,54]. Further details of this process, including a list of parameter estimates, can be found in the Supplementary Information.

**Figure 2.**
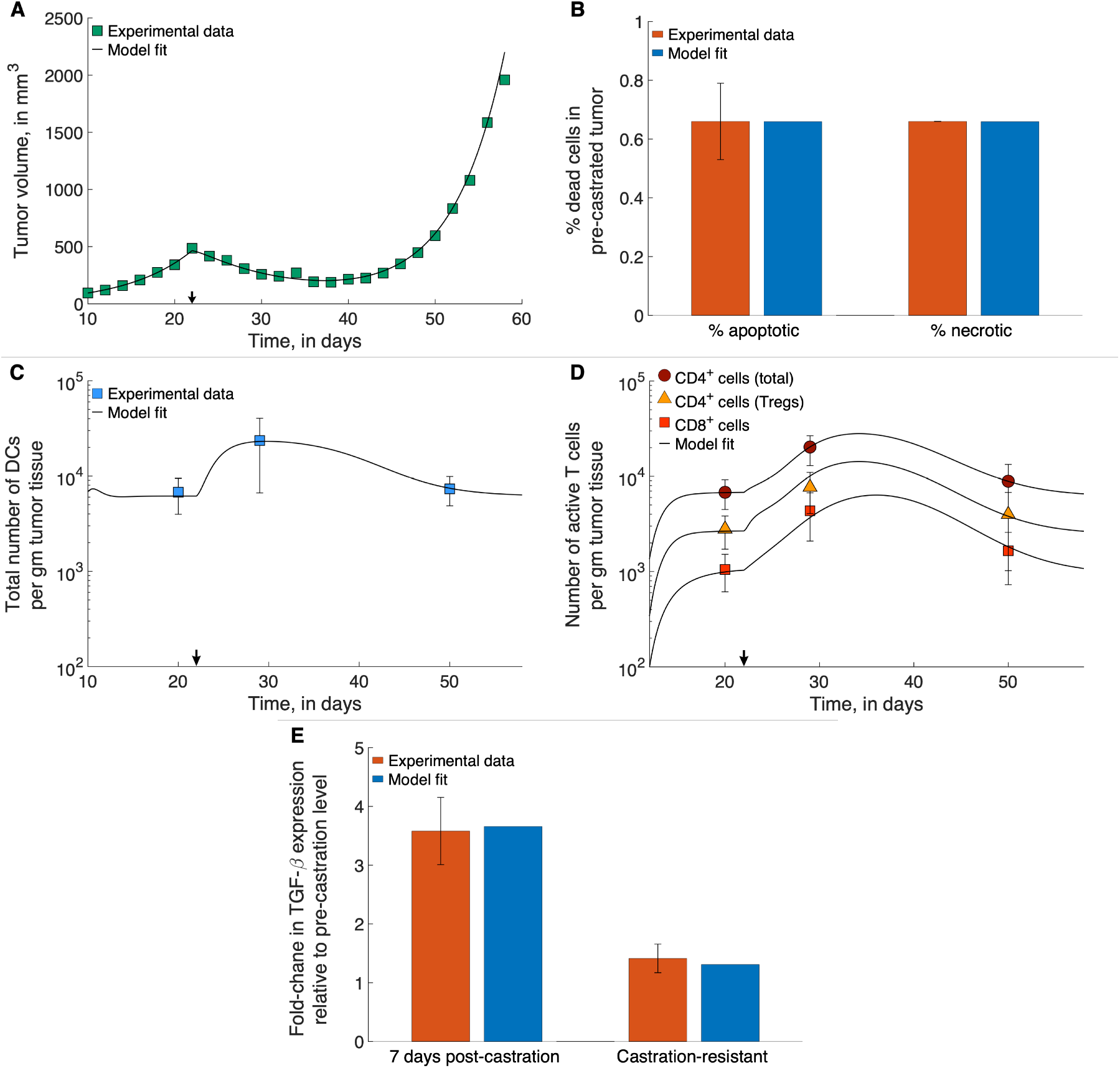
Model fits to data. (A), Model fitted to time-course treatment data on PCa xenograft volume, showing spontaneous emergence of castration resistance (black arrow indicates onset of ADT). (B), Model fitted to data on degree of apoptosis and necrosis in PCa xenografts, in the absence of treatment. (C), (D), Model fitted to time-course treatment data on PCa xenograft immune cell infiltration (black arrow indicates onset of ADT). (E), Model fitted to fold-changes in TGF-*β* expression within the xenografts 7-days post-ADT administration, and after the emergence of castration-resistance. Data were taken from [9,53,54].

### 2.3. The Standing Variations Method

Our model has 17 parameters that need to be estimated from the data shown in Figure 2. Biologically realistic values are assigned to a further 8 parameters, not characterized in the literature. We therefore expect a lack of practical identifiability or estimability in some of these parameters [55]. Furthermore, fixed parameter values, even if subject to uncertainty, would account only for standard error in experimental measurements, and be valid only for an ‘averaged’ individual mouse. Therefore, the fits shown in Figure 2 reflect samples from a distribution of parameter values which should reflect inter-mouse heterogeneity, and not experimental error. Capturing this heterogeneity is essential in understanding – and predicting – the response of a population to any therapeutic intervention.

Here, we propose the novel paradigm of *Standing Variations Modeling*, which exploits both, the uncertainty in data, and the inestimability of model parameters, to capture heterogeneity across a population with a deterministic model. Our method is outlined in Figure 3A. We begin by solving the inverse problem of inferring the probability distributions that model parameters follow, rather than estimating their precise values. Specifically, the multivariate uniform distribution is taken as a prior for the unknown model parameters. We then employ sampling importance resampling (SIR), a universally applicable method for obtaining draws from an unknown distribution [56], to infer posterior distributions on these parameters. Draws from the uniform priors are resampled based on their importance ratios, which measure the agreement between the approximated distribution and the experimental data, and are expected to be proportional to the resampling probabilities given the unknown distribution [57]. The SIR method has been successfully applied to quantify parameter uncertainty in epidemiology [58], systems pharmacology [59] and in nonlinear mixed effects models [57].

**Figure 3.**
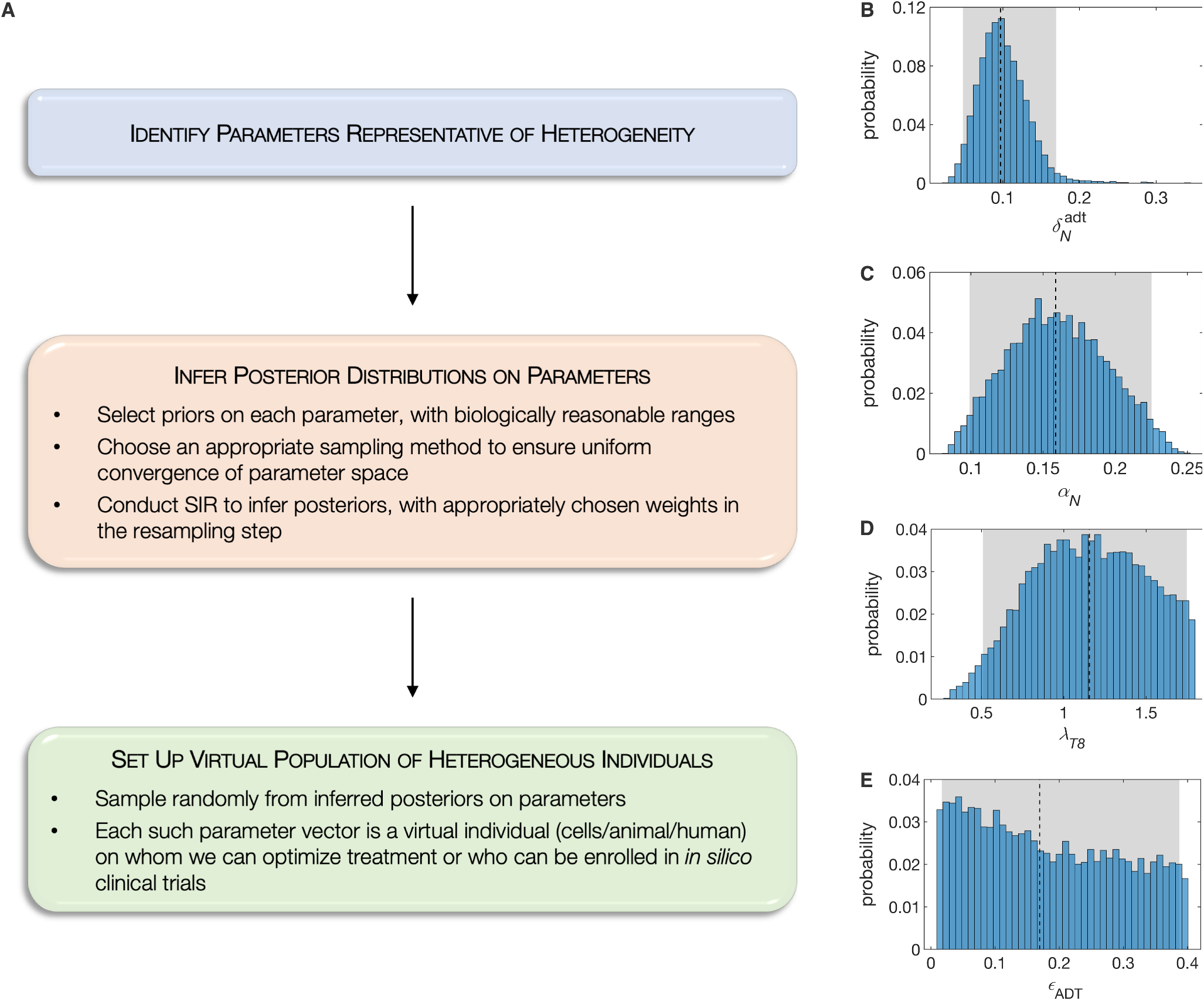
(A), The Standing Variations Modeling methodology. Representative inferred posterior distributions of model parameters, namely: (B), 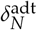, ADT-induced death rate of androgen sensitive cancer cells; (C), *α_N_*, proliferation rate of androgen sensitive cancer cell; (D), *λ_T_*_8_, maximum rate of naïve CD8^+^ T cell activation by mature APCs; and (E), *ϵ*_ADT_, fraction of ADT-induced cell death that is necrotic. Shown also are 95% confidence bounds (shaded gray areas) and median values (dashed black lines).

We selected 18 out of the 25 model parameters where identifiability issues are expected. This choice was motivated based on our biological knowledge of this system, since these parameters are likely to determine PCa response to ADT or immunotherapy. We first used Sobol sequences, a quasi Monte Carlo method [60,61], that allowed us to rapidly converge to a uniform sample from this high dimensional hypercube in parameter space. We then implemented the SIR method, assigning the original samples binary weights, based on whether or not they lie within the standard errors in experimental data points. The complete set of posterior distributions on these 18 parameters is available in the Supplementary Information. For illustration purposes, we plot four representative posterior distributions in Figure 3. As can be seen, the ADT-induced death rate of androgen sensitive cancer cells has narrow almost normal posterior, indicating that this parameter is, in fact, estimable from the given data (Figure 3B). On the other hand, the posterior on the necrotic fraction of ADT-induced cell death is almost uniform, indicating that this parameter is inestimable from the given data (Figure 3E).

In the final step of our Standing Variations Modeling approach, these posteriors on model parameters are sampled randomly to generate a collection of parameter vectors, each of whom is representative of a virtual individual. In our case, we generated a cohort of several thousand patients (mice) *in silico*, each characterized by their individualized parameters. An underlying assumption in this approach is that parameters that are inestimable from the given data, vary widely across the population. Having generated our population of test mice, we can conduct *in silico* clinical trials designed to predict the response of the population to sipuleucel-T, alone or in combination with ADT, as well as optimize the relative scheduling of these therapies.

### 2.4. In silico preclinical trial

For each preclinical trial simulated here, we first simulated a large enough cohort of virtual mice (as described above) so that we could randomize 5,000 mice in each treatment arm. The duration of these preclinical trials was fixed at 100 days. Following [9], the virtual animal was ‘euthanized’ or presumed dead when its tumor reached a size of 1500 mm^3^. We also assumed the animal was cured if its tumor volume shrank below 1 mm^2^, which is below typical detection thresholds. Overall survival was defined as the time period between treatment initiation and death.

### 2.5. Approximation of sensitivity analysis

To assess the contribution of model parameters to the distinct outcomes of a cure (tumor volume *<* 1 mm^3^) versus death (tumor volume exceed the survival threshold of 1500 mm^3^), we utilized a statistical approach: the two categories were fit as an outcome to a multiple logistic regression of the parameter values using the generalized linear model (in R, stats::glm) function in the statistical programming language and environment R. Because the relationship between probability of cure and parameter values may be non-linear, individually significant parameters were then fit to a second multiple logistic regression model as orthogonal polynomials (in R, stats::poly) of up to the third degree.

### 2.6. Treatment optimization

All treatment optimization was carried out on a cohort of 50,000 simulated mice, using a genetic algorithm [62]. Genetic algorithms mimic the principles of genetic evolution to solve optimization problems. For any desired objective function, such as maximizing the number of survivors or the median survival time, an initial pool of treatment strategies, for instance timing between doses, was created randomly. These strategies were then ranked based on the value of the objective function at the end of the treatment period. The top 20% of strategies were paired to create ‘offspring’ strategies. These inherit different properties from each of their parents and are also allowed a small chance of mutation. The resultant next generation of treatment protocols were sorted once again based on fitness, and the process repeated until convergence.

## 3. Results

### 3.1. Sipuleucel-T alone does not significantly improve overall survival of mice

We first investigated the effect on mouse survival, of the two treatments – ADT and vaccination with sipuleucel-T – administered as monotherapies. A cohort of 15,000 virtual mice was generated as described in Section 2.3. A preclinical trial was simulated as described in Section 2.4 with three treatment arms: control, ADT alone or vaccination alone. Our treatment protocols were consistent with those in [9]. That is, PCa xenografts were initiated in each mouse at day 0, and at day 22, when the xenografts reached an approximate volume of 400 mm^3^, the mice were randomly allocated to the treatment groups indicated. ADT was administered continuously to those in the ADT group and, similar to the clinical protocol for administering sipuleucel-T in humans, the animals in the vaccine group were inoculated weekly for a total of 3 doses.

The resultant Kaplan-Meier survival curves are plotted in Figure 4A. Median survival times for the mice in the control, vaccination, and ADT groups were 30 days, 32 days, and 58 days, respectively (p-values < 0.0001). The hazard ratio for death in the control group was 4.92 (95% CI, 4.67 - 5.19) compared to the ADT group, whilst it was 1.18 (95% CI, 1.14 - 1.23) compared to the vaccination group. Thus, our virtual mice population has a remarkably similar response to vaccination with sipuleucel-T, as the human population, with the vaccine conferring a very modest survival advantage over the control group. Our model suggests several potential reasons for this disappointing result. For instance, the inherent variability in the population, especially in terms of response to the vaccine, coupled with treatment initiation when the xenografts are already quite large (~ 400mm^3^ at day 22 post-implantation), and a strong immunosuppressive response from Tregs all contribute to the observations in Figure 4A.

**Figure 4.**
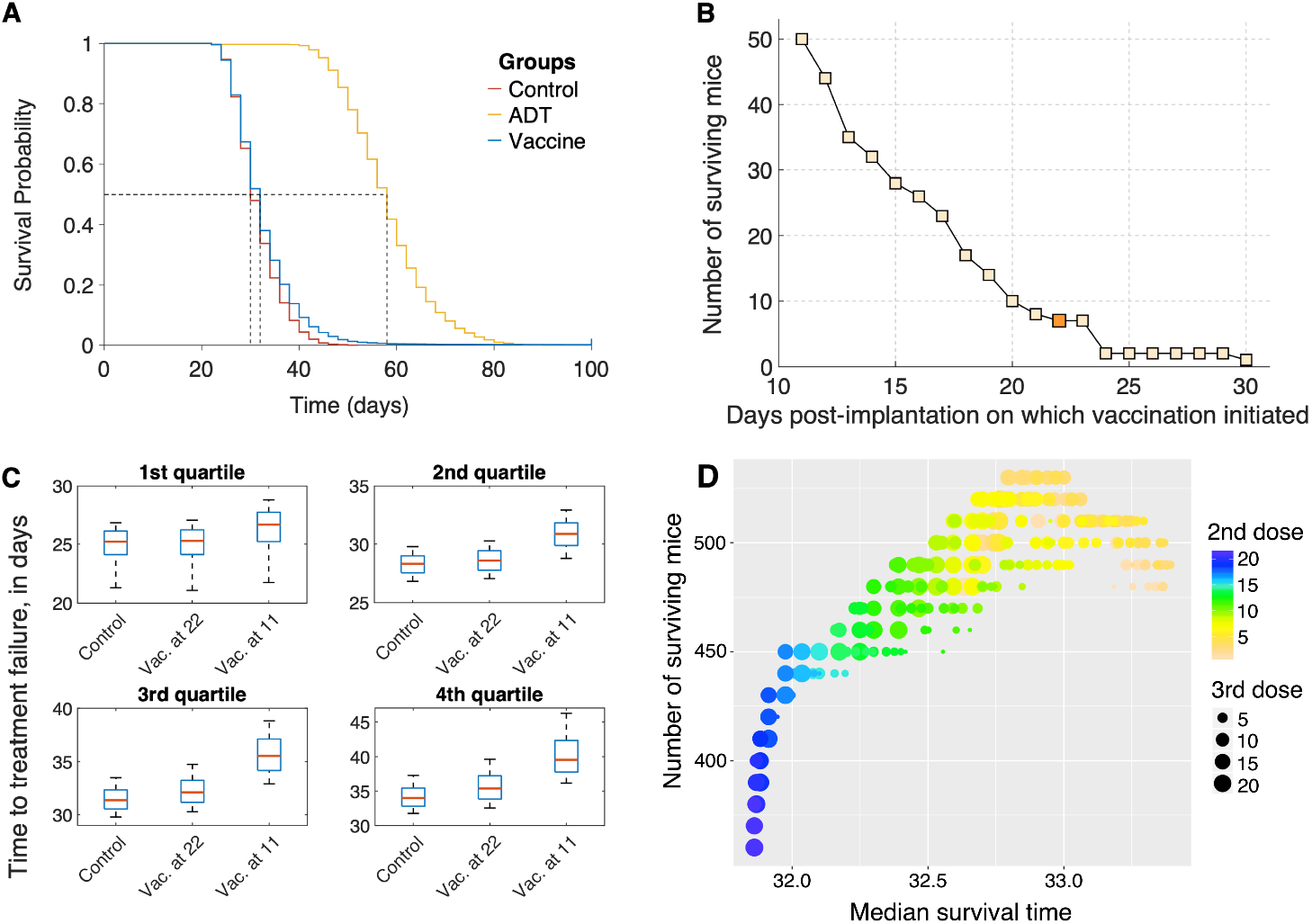
(A), Overall survival of simulated mice from time of therapy initiation. (B), Predicted number of mice that are alive at the end of the *in silico* trial (100 days), as a function of time post-xenograft implantation when first dose of sipuleucel-T was administered. Three total doses of vaccine were administered, given weekly. Dark shaded square corresponds to blue survival curve shown in panel (A). (C), Box plots of time to treatment failure (inter-quartile range and median) for the same simulated mice untreated (control), or recieving vaccination starting at either 22 or 11 days. Each subpanel corresponds to independent quartiles of the three strategies. (D), Optimization of sipuleucel-T scheduling. 3 total doses of vaccine were administered, with time of treatment initiation fixed at day 11 post-xenograft implantation, and times between second and third doses allowed to vary.

However, a closer inspection of the survival curve for mice on the vaccination arm (blue curve) reveals that 7 mice survived to the end of the preclinical trial. Although 7 mice out of a cohort of 5000 is small, this finding was a consistent prediction of the model across different cohorts of mice (data not shown). We therefore postulate that in rare cases, individuals may benefit significantly from sipuleucel-T.

### 3.2. Optimization of vaccine scheduling

We next investigated whether altering the schedule of vaccination could improve on the median overall survival and/or the number of surviving mice at the end of the trial. The results of this analysis are shown in Figure 4B-C.

Our model suggests that the survival benefit compared to control that vaccination confers is modest, in part, due to initiating treatment when the tumor is already large. We therefore varied the time of the first dose of vaccine on a fixed cohort of 5,000 mice. The earliest a dose could be given was at day 11 post-implantation, when the xenografts became palpable [9]. Clinically, this mimics starting vaccination at the time of first diagnosis. The remaining two doses were given weekly, as before. Figure 4B plots a graph of the number of surviving mice at the end of this trial, as a function of days post-implantation when vaccination was initiated. Starting treatment as early as possible confers a significant survival benefit to the population of mice, with the number of survivors increasing to 50 when vaccination was initiated on day 11 as compared to just 7 when vaccination was administered on day 22 (Figure 4B, dark shaded square). This result is explained by the fact that in our model, we account for a slow-down in the speed of CTL infiltration into the tumor with an increase in tumor volume [36].

Figure 4C shows the difference in survival/failure times for control mice, and for mice in vaccination schedules. The mice are divided into quartiles independently, to highlight heterogeneous benefits of early vaccination. Roughly 25% of mice (the first quartile) see no benefit from early vaccination. Another 25%, the fourth quartile, corresponding to mice with a relatively long survival even under control, show a 5 day increase in survival time for early vaccination. None of the simulated mice benefit appreciably from late vaccination, at 22 days.

An optimal dosing schedule has not been established for treating human patients with sipuleucel-T [8]. We therefore sought to maximize the median survival time by varying the intervals between successive doses of the vaccine. A cohort of 50,000 virtual mice was generated as described in Section 2.3. The timing of all three doses of vaccine were allowed to vary independently, with inter-dose times varying between 1 and 21 days. The earliest that treatment could be applied was at day 11 post-implantation (see comment in preceding paragraph). The optimal dosing schedule was identified by employing a genetic algorithm as described in Section 2.6. Unsurprisingly, initiating treatment at the earliest possible time-point (day 11) was optimal. Figure 4D plots the median survival time (*x*-axis) as a function of inter-dose intervals. The color of the dots indicates the interval between the first and second vaccine dose with blue dots corresponding to longer waiting times, while the size of the dots indicates the interval between the second and third vaccine dose with larger dots corresponding to longer waiting times. Median survival time was maximized when all three doses were administered 3-4 days apart. However, the median survival time only improved by 1.4 days, going from 32 to 33.4 days on this optimal protocol. These results offer an explanation as to why an optimal dosing schedule for sipuleucel-T has not been established in the clinic. Varying the inter-dose intervals across a large range of values only yielded a modest improvement in survival, which is possibly what has been observed in clinical trials of sipuleucel-T.

Encouraged by our finding that early administration of vaccine improves the survival chances of mice, we instead sought to maximize the number of survivors at the end of the trial, on the same cohort of 50,000 mice. Once again, initiating treatment at the earliest possible time-point (day 11) was found to be optimal. The number of surviving mice are plotted on the *y*-axis as a function of inter-dose intervals, in Figure 4D. A maximum of 580 mice (1.16%) survive, of which 395 are cured (tumor size *<* 1 mm^3^), if the first two vaccine doses are given within 3 days of each other, whilst the third dose is administered 13-15 days later. This is a marked improvement from the 7 survivors out of 5,000 (0.14%) when mice were vaccinated starting on day 22, on a weekly schedule. We remark that the number of surviving mice (equivalently, the higher quantiles of the survival time) is more sensitive to differences in strategy than is the median survival time. This is typical in biomedical studies where only a subpopulation of patients benefit from the intervention (for a review, see [63]).

### 3.3. Characteristics of response to vaccine

The characteristics of the surviving mice – and in particular, the rare, cured subpopulation – may be of crucial importance in determining who will benefit the most from vaccination, and to what extent. The power of a mechanistic model is to extrapolate into these desirable regions of parameter space which can then be targeted and explored in a clinical setting.

We therefore sought to identify those parameters that are significantly associated with a cure (tumor size *<* 1 mm^3^). Table 1 gives trends in model parameters significantly associated with this outcome in an individual simulated mouse, from the final cohort of 50,000 mice. Multiple logistic regression compared 395 cured mice to 49,502 ‘euthanized’ or dead mice, with mice surviving-but-not-cured at the end of the simulation excluded from this analysis. Positive effect sizes indicate terms of polynomials of parameters that are higher in the cured mice, while negative effect sizes indicate the opposite. The p-values can be made arbitrarily small by increasing the number of simulated mice, but the relative value of Z-scores between parameters reflects their contribution to the outcome, approximating a sensitivity analysis if the model could be analyzed in such a way. Non-significant parameters are not reported in the table, for the sake of brevity.

**Table 1.**
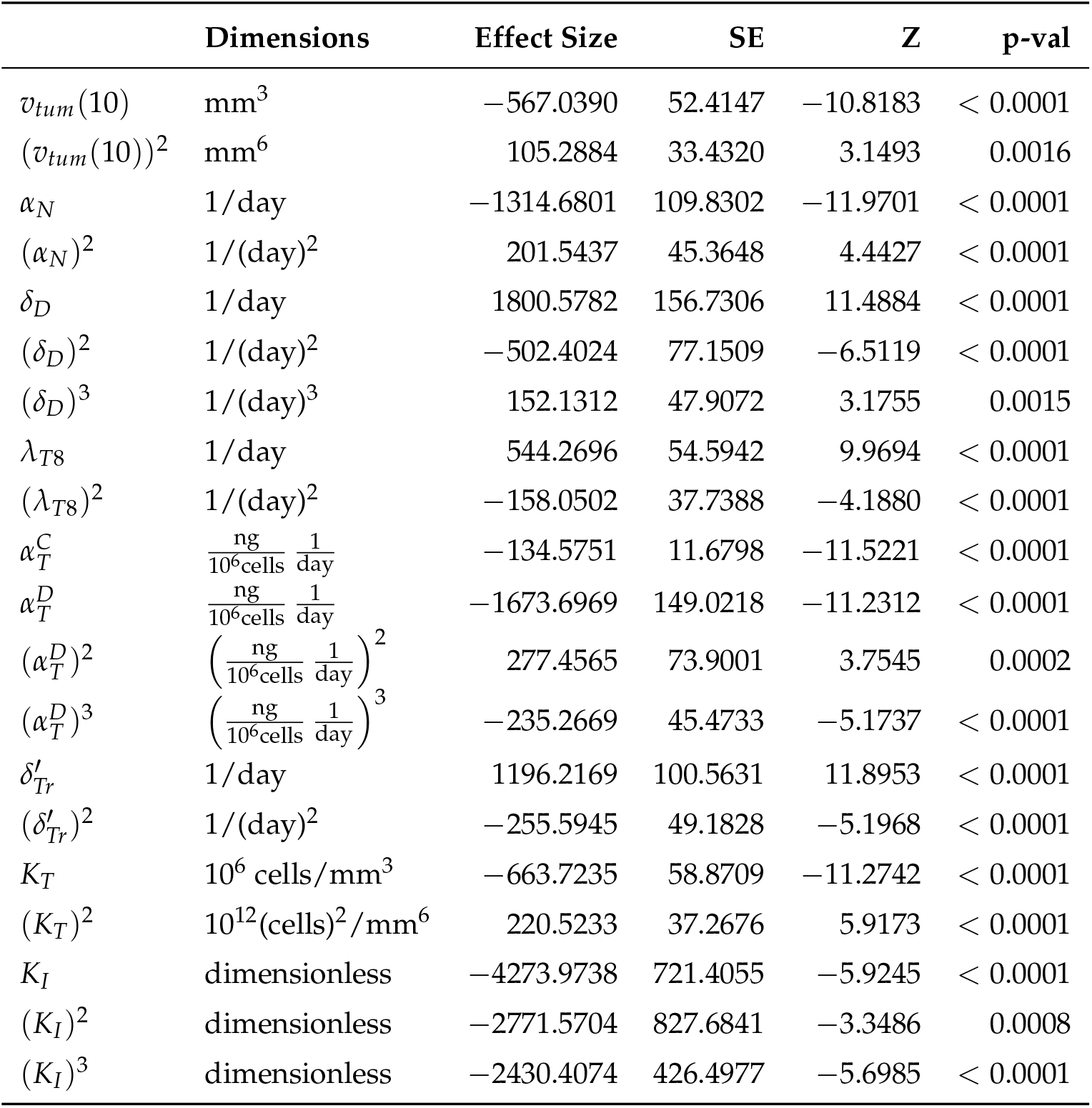
For each parameter, including higher-order terms where significant: the Dimensions of the parameter, the Effect Size (on a logistic scale, per unit), the standard error in the estimate of the effect size (SE), the corresponding Z score and p-value from the multiple logistic regression.

|  | Dimensions | Effect Size | SE | Z | p-val |
| --- | --- | --- | --- | --- | --- |
| $v_{tum}(10)$ | mm<sup>3</sup> | −567.0390 | 52.4147 | −10.8183 | < 0.0001 |
| $(v_{tum}(10))^2$ | mm<sup>6</sup> | 105.2884 | 33.4320 | 3.1493 | 0.0016 |
| $\alpha_N$ | 1/day | −1314.6801 | 109.8302 | −11.9701 | < 0.0001 |
| $(\alpha_N)^2$ | 1/(day)<sup>2</sup> | 201.5437 | 45.3648 | 4.4427 | < 0.0001 |
| $\delta_D$ | 1/day | 1800.5782 | 156.7306 | 11.4884 | < 0.0001 |
| $(\delta_D)^2$ | 1/(day)<sup>2</sup> | −502.4024 | 77.1509 | −6.5119 | < 0.0001 |
| $(\delta_D)^3$ | 1/(day)<sup>3</sup> | 152.1312 | 47.9072 | 3.1755 | 0.0015 |
| $\lambda_{T8}$ | 1/day | 544.2696 | 54.5942 | 9.9694 | < 0.0001 |
| $(\lambda_{T8})^2$ | 1/(day)<sup>2</sup> | −158.0502 | 37.7388 | −4.1880 | < 0.0001 |
| $\alpha_T^C$ | $\frac{\text{ng}}{10^6 \text{cells}} \frac{1}{\text{day}}$ | −134.5751 | 11.6798 | −11.5221 | < 0.0001 |
| $\alpha_T^D$ | $\frac{\text{ng}}{10^6 \text{cells}} \frac{1}{\text{day}}$ | −1673.6969 | 149.0218 | −11.2312 | < 0.0001 |
| $(\alpha_T^D)^2$ | $\left(\frac{\text{ng}}{10^6 \text{cells}} \frac{1}{\text{day}}\right)^2$ | 277.4565 | 73.9001 | 3.7545 | 0.0002 |
| $(\alpha_T^D)^3$ | $\left(\frac{\text{ng}}{10^6 \text{cells}} \frac{1}{\text{day}}\right)^3$ | −235.2669 | 45.4733 | −5.1737 | < 0.0001 |
| $\delta'_{Tr}$ | 1/day | 1196.2169 | 100.5631 | 11.8953 | < 0.0001 |
| $(\delta'_{Tr})^2$ | 1/(day)<sup>2</sup> | −255.5945 | 49.1828 | −5.1968 | < 0.0001 |
| $K_T$ | 10<sup>6</sup> cells/mm<sup>3</sup> | −663.7235 | 58.8709 | −11.2742 | < 0.0001 |
| $(K_T)^2$ | 10<sup>12</sup>(cells)<sup>2</sup>/mm<sup>6</sup> | 220.5233 | 37.2676 | 5.9173 | < 0.0001 |
| $K_I$ | dimensionless | −4273.9738 | 721.4055 | −5.9245 | < 0.0001 |
| $(K_I)^2$ | dimensionless | −2771.5704 | 827.6841 | −3.3486 | 0.0008 |
| $(K_I)^3$ | dimensionless | −2430.4074 | 426.4977 | −5.6985 | < 0.0001 |

<sup>1</sup> Since we are administering vaccination alone, we do not distinguish between an androgen-sensitive versus castration-resistant phenotype.

The significant parameters were found to be: *v_tum_*(10), the xenograft tumor volume at day 10 (1 day prior to vaccination initiation); *α_N_*, the rate of cancer cell proliferation^1^; *δ_D_*, the rate of dead cell clearance by (assumed) macrophages; *λ_T_*_8_, the maximum rate of naïve CTL activation by mature APCs; 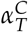, the rate of TGF-*β* expression by cancer cells; 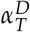, the rate of TGF-*β* expression by (assumed) macrophages, taken here to be proportional to the number of dead cells; 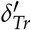, the rate of activated Treg death; *K_T_*, the half-saturation constant for APC-mediated activation of T cells; and *K_I_*, the CTL to target cell ratio at which rate of CTL-induced cell kill is half its maximum value. For most parameters, first and second order terms have opposite effects, indicating “diminishing returns”; exceptions being 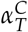, which shows a linear relationship, and *K_I_*, which shows a compounding effect. The overall differences in parameter values are summarized as box plots in Figure 5.

**Figure 5.**
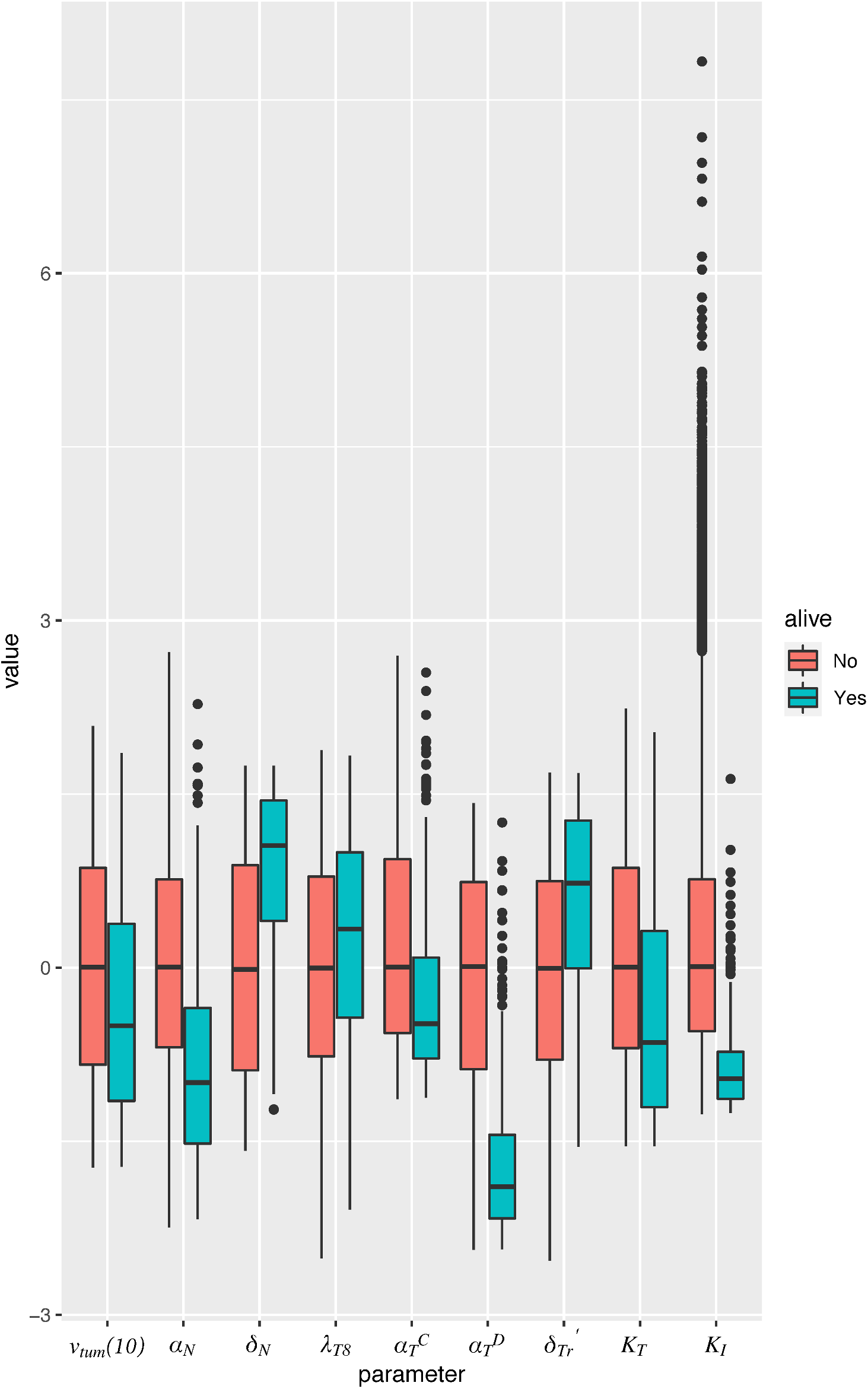
Box plot of parameter values in simulated mice, giving median and interquartile range. Parameters are normalized to mean and standard deviation across all simulated mice. Only those parameters are represented that were found to be significantly associated with a cure (tumor size 1 mm^3^) when vaccination is administered as a monotherapy, on the optimal schedule predicted by the model. Red boxes correspond to dead mice, and teal boxes correspond to cured mice.

From the Z-scores in Table 1, we conclude that the most important determinants of response to vaccination are *α_N_* and 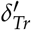, with a low tumor cell proliferation rate or a high rate of activated Treg death associated with cured mice. The next largest effects come from *δ_D_*, 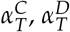 and *K_T_*, with high values of *δ_D_* and low values of 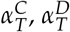 and *K_T_* associated with cured mice. *δ_D_*, 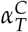 and 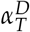 characterize the amount of TGF-*β* in the system: a large *δ_D_* would result in more efficient clearance of dead cells from the tumor space, and thus correspond to reduced macrophage numbers, and a smaller source of TGF-*β*. Equally, small 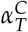 or 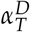 would indicate diminished TGF-*β* production and consequently, elevated CTL activity.

### 3.4. ADT and vaccination combination therapy

We investigated the potential of combining sipuleucel-T with ADT, and sought an optimal relative dosing schedule that maximizes median overall survival. As before, optimization was carried out on a cohort of 50,000 simulated mice, with timing of ADT initiation fixed at day 21 post-implantation. Three doses of vaccine could be administered with inter-dose timing allowed to vary between 1 and 21 days. The earliest that vaccination could be started was at day 11 post-implantation. The optimal dosing schedule, identified by employing a genetic algorithm, was vaccine doses at days 12, 16 and 27 post-implantation. That is, our model suggests administering two doses of vaccine in quick succession, and waiting for the third dose until after ADT initiation.

To illustrate the potential benefit of combining vaccination with ADT across a population, we conducted an *in silico* preclinical trial, comparing: (1) our predicted optimal dose schedule; (2) initiating vaccination simultaneously with ADT (vaccine given in three weekly doses); and (3) administering vaccination 7 days post-ADT, when castration resistance is observed in the mice xenografts [9]. We remark that this last schedule mimics the current clinical protocol for administering sipuleucel-T. As before, 15,000 virtual mice enrolled in our trial and 5,000 mice randomized into each treatment arm. The resultant Kaplan-Meier survival curves are plotted in Figure 6A. Median survival times for the mice in the clinical arm (schedule (3)), simultaneous arm (schedule (2)), and optimal arm (schedule (1)) were 58 days, 62 days, and 64 days, respectively (p-values < 0.0001). The hazard ratio in the clinical group was 1.58 (95% CI, 1.52 - 1.65) compared to the simultaneous group, whilst it was 1.79 (95% CI, 1.72 - 1.87) compared to the optimal schedule group. Moreover, only 20 mice (0.4%) were predicted to survive to he end of the trial on the clinical protocol. When sipuleucel-T was co-administered with ADT, there were 284 survivors (~5.7%) with 179 cured, whilst this number went up to 441 (~8.8%) with 318 cured on the optimal protocol. Thus, our model suggests that instead of waiting until after ADT has failed, vaccination should be given *prior* to starting androgen deprivation, and in a staggered, rather than periodic fashion. This is predicted to reduce the rate of cancer death by approximately 45%.

**Figure 6.**
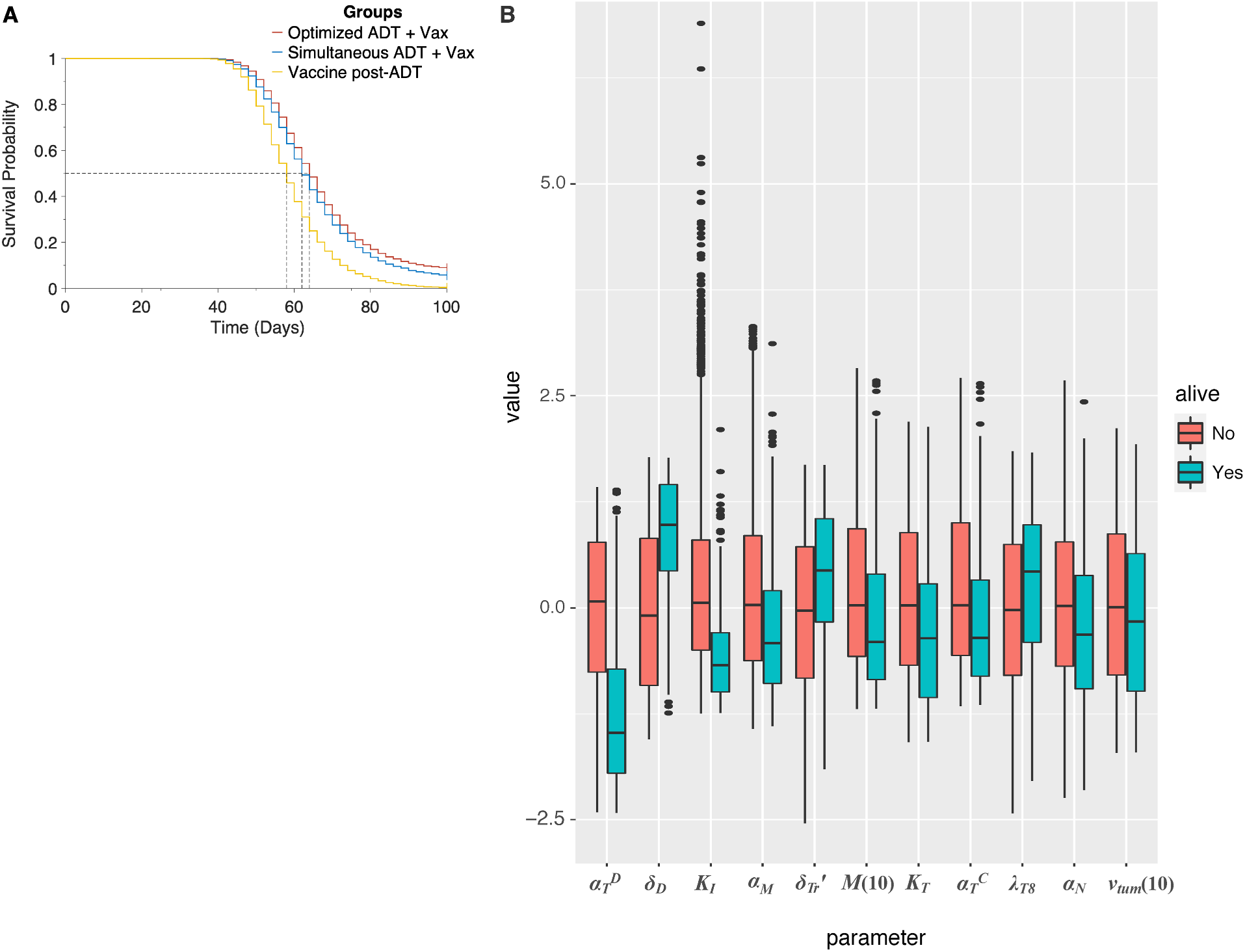
(A), Overall survival of simulated mice from time of combination treatment initiation, on various schedules. (B), Box plot of parameter values in simulated mice, giving median and inter-quartile range. Parameters are normalized to mean and standard deviation across all simulated mice. Only those parameters are represented that were found to be significantly associated with a cure (tumor size 1 mm^3^) when ADT and vaccination are administered as a combination, on the optimal schedule predicted by the model. Red boxes correspond to dead mice, and teal boxes correspond to cured mice.

Finally, we sought to identify those parameters that are significantly associated with a cure (tumor size *<* 1 mm^3^), when ADT is combined with vaccination and administered per the optimal schedule found above. In addition to the parameters found to be significantly associated with a cure when vaccination is administered as a monotherapy (see Section 3.3), now *α_M_*, the rate of castration-resistant cell proliferation, and *M*(10), the number of castration resistant cells that are present in the tumor prior to treatment initiation are also predicted to be significant. The overall differences in parameter values are summarized as box plots in Figure 6B (see also Table S3.1 in the Supplementary Information).

To illustrate the potential benefit of combining vaccination with ADT across a population, we conducted an *in silico* preclinical trial, comparing: (1) our predicted optimal dose schedule; (2) initiation of vaccination simultaneously with ADT (vaccine given in three weekly doses); and (3) administration of vaccination 7 days post-ADT, when castration resistance is observed in the mice xenografts [9]. This last schedule mimics the current clinical protocol for sipuleucel-T. As before, 15,000 virtual mice enrolled in our trial and 5,000 mice were randomized into each treatment arm. The resultant Kaplan-Meier survival curves are shown in Figure 6A. Median survival times for the mice in the clinical arm (schedule (3)), simultaneous arm (schedule (2)), and optimal arm (schedule (1)) were 58 days, 62 days, and 64 days, respectively (p-values < 0.0001). The hazard ratio in the clinical group was 1.58 (95% CI, 1.52 - 1.65) compared to the simultaneous arm and 1.79 (95% CI, 1.72 - 1.87) compared to the optimal schedule arm. Moreover, only 20 mice (0.4%) were predicted to survive to the end of the trial in arm 3. When sipuleucel-T was co-administered with ADT, there were 284 survivors (~5.7%) with 179 cured, and 441 survivors (~8.8%) with 318 cured on the optimal protocol. Thus, our model suggests that instead of waiting until after ADT has failed, vaccination should be given *prior* to starting androgen deprivation, and in a staggered, rather than periodic fashion. This is predicted to reduce the rate of cancer death by approximately 45%.

Finally, we sought to identify those parameters that are significantly associated with a cure (tumor size *<* 1 mm^3^), when ADT is combined with vaccination and administered per the optimal schedule found above. In addition to the parameters found to be significantly associated with a cure when vaccination is administered as a monotherapy (see Section 3.3), now *α_M_*, the rate of castration-resistant cell proliferation, and *M*(10), the number of castration resistant cells that are present in the tumor prior to treatment initiation are also predicted to be significant. The overall differences in parameter values are summarized as box plots in Figure 6B (see also Table S3.1 in the Supplementary Information).

## 4. Discussion

The landscape for PCa therapy has developed rapidly with the approval of new classes of treatments, including immune-based approaches, for use in various PCa disease states [4]. FDA approval of sipuleucel-T for the treatment of castration resistant PCa marked the first cancer “vaccine” approved for use in a treatment setting and sparked renewed interest in cancer vaccines more broadly [64]. Despite the initial enthusiasm for sipuleucel-T, it has not been widely used compared with other available therapies, in part, due to its limited clinical success [7,65].

A powerful and practical way to analyze potential mechanisms driving therapeutic success and failure, and to optimize dosing schedules of novel drugs, is to use multi-scale computational (dynamical systems) modeling of cancer response to treatment. Biomedically sound parametrization is essential to the validity of such modeling approaches. Validated mechanistic models enable reliable, quantitative predictions [55,66]. However, data are typically sparse or incomplete, leading to parameter identifiability issues [55]. Furthermore, even if some parameter values can be determined precisely, these would only be valid for an ‘averaged’ individual (cell/mouse/human), and cannot capture the inherent variability across a population, let alone within an individual.

In this paper we developed a detailed and integrative mathematical model of PCa response to sipuleucel-T and ADT, in order to shed light on the limited clinical success of the sipuleucel-T vaccine. Our model simulates the growth and response to treatment of PCa xenografts in an *individual* (mouse). However, we aim to understand the response of a *population* to vaccine therapy. Therefore, we proposed a novel modeling paradigm that exploits uncertainty and variability in data to capture heterogeneity across a population. This heterogeneity is key to understanding both health disparities in PCa outcomes (including especially ethnic/racial disparities [2]), and the results sipuleucel-T clinical trials. We term this synthesis of importance sampling and deterministic differential equation modeling a ‘Standing Variations Model’, inspired by similar approaches in population genetics [10,11].

Using our modeling framework, we first conducted an *in silico* preclinical trial of sipuleucel-T administered as a monotherapy. Survival data from our simulated cohort of mice was found to be in good qualitative agreement with results from the first phase III clinical trial of sipuleucel-T [7], with vaccination providing only a modest survival benefit compared to control. In our model, several parameters that are key determinants of response to vaccination vary significantly across the population, and no one parameter in and of itself determines a favorable outcome. However, these parameter differences do suggest mechanistic causes of limited sipuleucel-T efficacy, in both our simulated mice and in the clinic. For example, tumor sizes are likely too large at the time of treatment initiation, making it almost impossible for active CTLs to successfully infiltrate the tumor and eliminate target cancer cells. Further, the vaccine elicits a strong immunosuppressive response in terms of Treg activation and TGF-*β* expression within the tumor, both of which appear to limit efficacy.

We next varied vaccination schedules to: (1) understand why an optimal dosing schedule has not been established for sipuleucel-T [8]; and (2) what such a schedule would be. Our simulations indicated that even varying the time interval between vaccine doses over a wide range could not appreciably improve on median overall survival of the mouse population. This is again in agreement with what has been observed clinically for sipuleucel-T [8]. However, a small fraction of the population was predicted to benefit significantly from vaccination, motivating us to instead seek treatment protocols that maximize the number of such individuals. A non-standard dosing schedule – two quick doses followed by a longer interval to the third dose – was predicted to be optimal. These results support an off-label trial, with investigational use of sipuleucel-T in patients with androgen-sensitive cancer to rigorously assess potential survival benefits.

Sipuleucel-T is currently approved for the treatment of patients with mCRPC, that is, after ADT has failed [4]. We therefore conducted a preclinical trial to investigate the potential of combining ADT with vaccination. Our model predicted administering 2 doses of the vaccine *prior* to ADT was optimal, in terms of maximizing the median overall survival, and resulted in 6.36% cured mice as compared to 0.08% on the schedule that mimics the current clinical protocol. This suggests that there is potential to significantly improve the clinical performance of sipuleucel-T by combining it with other therapeutic modalities such as ADT. Furthermore, a multiple logistic regression analysis was conducted to identify the parameters that are significantly associated with a cure (tumor volume *<* 1 mm^3^) under vaccination alone, or vaccination in combination with ADT. Parameters associated with a substantial immunosuppressive response to the vaccine, such as TGF-*β* production rates, were strongly negatively correlated with this favorable outcome. These results highlight a potential for combining anti-TGF-*β* therapy with vaccination in improving outcomes, although this would require investigational agents such as Galunisertib [67].

The standing variation modeling approach introduced here is flexible enough to include these agents, as well as to model adaptive clinical trial designs incorporating any of these parameters as biomarkers. Work is ongoing on future iterations of this model which will include additional combination therapies and design elements, as well as additional mechanistic features such as the activation or differentiation of monocyte lineage cells (for instance, macrophages) and incorporating immunosuppressive functions of cancer cells via the PD1-PDL1 axis [68].

Clinical translation of biological insights, such as those underlying cancer vaccines, requires tremendous investments of time, money, and limited clinical-scientific resources such as trained physician-scientists and the recruitment of rare patient populations. Mathematical modeling can both, predict the success of these investments, and optimize that success given that even safe and effective novel therapeutic modalities may provide limited clinical benefit if administered in a sub-optimal way. This optimization is a particular strength of our approach, since clinical trials randomizing hundreds of thousands of mice or patients to different combination therapies, each administered over a range of potential schedules, would be prohibitively expensive.

## Author Contributions

Conceptualization, H.V.J. and T.L.J.; methodology, H.V.J.; software, H.V.J., I.C.S. and S.K.H.; formal analysis, H.V.J. and S.K.H.; investigation, H.V.J., I.C.S., S.K.H., J.B. and T.L.J.; writing—original draft preparation, H.V.J., S.K.H. and T.L.J.; writing—review and editing, H.V.J., I.C.S., S.K.H., J.B. and T.L.J.; visualization, H.J, I.C.S. and S.K.H.; supervision, H.V.J. and T.L.J. All authors have read and agreed to the published version of the manuscript.

## Funding

This research received no external funding.

## Institutional Review Board Statement

Not applicable

## Informed Consent Statement

Not applicable

## Acknowledgments

The authors thank Ms. Krishna Natarajan and Prof Giray Okten for many helpful discussions.

## Conflicts of Interest

The authors declare no conflict of interest.

## Abbreviations

The following abbreviations are used in this manuscript:

APC: Antigen-presenting cell
CTL: CD8^+^ Cytotoxic T lymphocyte DC Dendritic cell
mCRPC: Metastatic castration-resistant prostate cancer
ODE: Ordinary differential equation
PAP: Prostatic acid phosphatase
PBMC: Peripheral blood mononuclear cell
PCa: Prostate cancer
Th cell: CD4^+^ T helper cell
Treg: CD4^+^ regulatory T cell

## Footnotes

1 Since we are administering vaccination alone, we do not distinguish between an androgen-sensitive versus castration-resistant phenotype.

## Notes

### Competing Interest Statement

The authors have declared no competing interest.

## References

1. Siegel, R.L.; Miller, K.D.; Jemal, A. Cancer statistics, 2020. CA: A Cancer Journal for Clinicians 2020, 70, 7–30.

2. Lewis, D.D.; Cropp, C.D. The impact of African ancestry on prostate cancer disparities in the era of precision medicine. Genes 2020, 11, 1471.

3. Perlmutter, M.A.; Lepor, H. Androgen deprivation therapy in the treatment of advanced prostate cancer. Reviews in Urology 2007, 9, S3–S8.

4. Bilusic, M.; Madan, R.A.; Gulley, J.L. Immunotherapy of prostate cancer: facts and hopes. Clinical Cancer Research 2017, 23, 6764–6770.

5. Komura, K.; Sweeney, C.J.; Inamoto, T.; Ibuki, N.; Azuma, H.; Kantoff, P.W. Current treatment strategies for advanced prostate cancer. International Journal of Urology 2018, 25, 220–231.

6. Han, P.; Hanlon, D.; Sobolev, O.; Chaudhury, R.; Edelson, R.L. Ex vivo dendritic cell generation – A critical comparison of current approaches. In International Review of Cell and Molecular Biology; Lhuillier, C.; Galluzzi, L., Eds.; Academic Press, 2019; Vol. 349, pp. 251–307.

7. Kantoff, P.W.; Higano, C.S.; Shore, N.D.; Berger, E.R.; Small, E.J.; Penson, D.F.; Redfern, C.H.; Ferrari, A.C.; Dreicer, R.; Sims, R.B.; others. Sipuleucel-T immunotherapy for castration-resistant prostate cancer. New England Journal of Medicine 2010, 363, 411–422.

8. Provenge – FDA. Available online: https://www.fda.gov/media/78511/download (accessed January 05, 2020).

9. Shen, Y.C.; Ghasemzadeh, A.; Kochel, C.M.; Nirschl, T.R.; Francica, B.J.; Lopez-Bujanda, Z.A.; Haro, M.A.C.; Tam, A.; Anders, R.A.; Selby, M.J.; Korman, A.J.; Drake, C.G. Combining intratumoral Treg depletion with androgen deprivation therapy (ADT): preclinical activity in the Myc-CaP model. Prostate Cancer and Prostatic Diseases 2018, 21, 113–125.

10. Birdsell, J.B. Evolution, genetics, and man. By Theodosius Dobzhansky, 1955. John Wiley and Sons, New York, IX, 398 pp., $5.50. American Journal of Physical Anthropology 1956, 14, 665–668.

11. Nei, M. Genetic polymorphism and neomutationism. Evolutionary Dynamics of Genetic Diversity; Mani, G.S., Ed.; Springer: Berlin, Heidelberg, 1984; pp. 214–241.

12. Ideta, A.M.; Tanaka, G.; Takeuchi, T.; Aihara, K. A mathematical model of intermittent androgen suppression for prostate cancer. Journal of Nonlinear Science 2008, 18, 593–614.

13. Jackson, T.L. A mathematical model of prostate tumor growth and androgen-independent relapse. Discrete and Continuous Dynamical Systems - Series B 2004, 4, 187–201.

14. Jackson, T.L. A mathematical investigation of the multiple pathways to recurrent prostate cancer: comparison with experimental data. Neoplasia 2004, 6, 697–704.

15. Kronik, N.; Kogan, Y.; Elishmereni, M.; Halevi-Tobias, K.; Vuk-Pavlović, S.; Agur, Z. Predicting outcomes of prostate cancer immunotherapy by personalized mathematical models. Plos One 2010, 5, e15482.

16. Tanaka, G.; Hirata, Y.; Goldenberg, S.; Bruchovsky, N.; Aihara, K. Mathematical modelling of prostate cancer growth and its application to hormone therapy. Philosophical Transactions of the Royal Society of London A: Mathematical, Physical and Engineering Sciences 2010, 368, 5029–5044.

17. Jain, H.V.; Clinton, S.K.; Bhinder, A.; Friedman, A. Mathematical modeling of prostate cancer progression in response to androgen ablation therapy. Proceedings of the National Academy of Sciences 2011, 108, 19701–19706.

18. Portz, T.; Kuang, Y.; Nagy, J.D. A clinical data validated mathematical model of prostate cancer growth under intermittent androgen suppression therapy. AIP Advances 2012, 2, 011002.

19. Jain, H.V.; Friedman, A. Modeling prostate cancer response to continuous versus intermittent androgen ablation therapy. Discrete and Continuous Dynamical Systems - Series B 2013, 18, 945–967.

20. Morken, J.D.; Packer, A.; Everett, R.A.; Nagy, J.D.; Kuang, Y. Mechanisms of resistance to intermittent androgen deprivation in patients with prostate cancer identified by a novel computational method. Cancer Research 2014, 74, 3673–83.

21. Brady-Nicholls, R.; Nagy, J.D.; Gerke, T.A.; Zhang, T.; Wang, A.Z.; Zhang, J.; Gatenby, R.A.; Enderling, H. Prostate-specific antigen dynamics predict individual responses to intermittent androgen deprivation. Nature Communications 2020, 11, 1–13.

22. Mahlbacher, G.E.; Reihmer, K.C.; Frieboes, H.B. Mathematical modeling of tumor-immune cell interactions. Journal of Theoretical Biology 2019, 469, 47–60.

23. Wilkie, K.P. A Review of Mathematical Models of Cancer–Immune Interactions in the Context of Tumor Dormancy. In Systems Biology of Tumor Dormancy; Enderling, H.; Almog, N.; Hlatky, L., Eds.; Springer: New York, NY, 2013; pp. 201–234.

24. Bądziul, D.; Jakubczyk, P.; Chotorlishvili, L.; Toklikishvilie, Z.; Traciak, J.; Jakubowicz-Gil, J.; Chmiel-Szajner, S. Mathematical prostate cancer evolution: effect of immunotherapy based on controlled vaccination strategy. Computational and Mathematical Methods in Medicine 2020, 2020, 7970265.

25. Coletti, R.; Leonardelli, L.; Parolo, S.; Marchetti, L. A QSP model of prostate cancer immunotherapy to identify effective combination therapies. Scientific reports 2020, 10, 1–18.

26. Rutter, E.M.; Kuang, Y. Global dynamics of a model of joint hormone treatment with dendritic cell vaccine for prostate cancer. Discrete and Continuous Dynamical Systems - Series B 2017, 22, 1001–1021.

27. Radunskaya, A.; Kim, R.; Woods II, T. Mathematical modeling of tumor immune interactions: a closer look at the role of a PD-L1 inhibitor in cancer immunotherapy. Spora: A Journal of Biomathematics 2018, 4, 25–41.

28. Robertson-Tessi, M.; El-Kareh, A.; Goriely, A. A mathematical model of tumor–immune interactions. Journal of theoretical biology 2012, 294, 56–73.

29. Fink, S.L.; Cookson, B.T. Apoptosis, pyroptosis, and necrosis: mechanistic description of dead and dying eukaryotic cells. Infection and Immunity 2005, 73, 1907–1916.

30. Green, D.; Oguin, T.; Martinez, J. The clearance of dying cells: table for two. Cell Death and Differentiation 2016, 23, 915–926.

31. Elliott, M.R.; Ravichandran, K.S. The dynamics of apoptotic cell clearance. Developmental Cell 2016, 38, 147–160.

32. Sauter, B.; Albert, M.L.; Francisco, L.; Larsson, M.; Somersan, S.; Bhardwaj, N. Consequences of cell death: exposure to necrotic tumor cells, but not primary tissue cells or apoptotic cells, induces the maturation of immunostimulatory dendritic cells. Journal of Experimental Medicine 2000, 191, 423–434.

33. Gardner, A.; Ruffell, B. Dendritic cells and cancer immunity. Trends in Immunology 2016, 37, 855–865.

34. Banchereau, J.; Steinman, R.M. Dendritic cells and the control of immunity. Nature 1998, 392, 245–252.

35. Mitra, S.; Leonard, W.J. Biology of IL-2 and its therapeutic modulation: mechanisms and strategies. Journal of Leukocyte Biology 2018, 103, 643–655.

36. Boissonnas, A.; Fetler, L.; Zeelenberg, I.S.; Hugues, S.; Amigorena, S. In vivo imaging of cytotoxic T cell infiltration and elimination of a solid tumor. Journal of Experimental Medicine 2007, 204, 345–356.

37. Martínez-Lostao, L.; Anel, A.; Pardo, J. How do cytotoxic lymphocytes kill cancer cells? Clinical Cancer Research 2015, 21, 5047–5056.

38. Wing, J.B.; Tanaka, A.; Sakaguchi, S. Human FOXP3+ regulatory T cell heterogeneity and function in autoimmunity and cancer. Immunity 2019, 50, 302–316.

39. Miyara, M.; Yoshioka, Y.; Kitoh, A.; Shima, T.; Wing, K.; Niwa, A.; Parizot, C.; Taflin, C.; Heike, T.; Valeyre, D.; others. Functional delineation and differentiation dynamics of human CD4+ T cells expressing the FoxP3 transcription factor. Immunity 2009, 30, 899–911.

40. Yamazaki, S.; Iyoda, T.; Tarbell, K.; Olson, K.; Velinzon, K.; Inaba, K.; Steinman, R.M. Direct expansion of functional CD25+ CD4+ regulatory T cells by antigen-processing dendritic cells. Journal of Experimental Medicine 2003, 198, 235–247.

41. Yadav, M.; Bluestone, J.A.; Stephan, S. Peripherally induced tregs–role in immune homeostasis and autoimmunity. Frontiers in Immunology 2013, 4, 232.

42. Alonso, R.; Flament, H.; Lemoine, S.; Sedlik, C.; Bottasso, E.; Péguillet, I.; Prémel, V.; Denizeau, J.; Salou, M.; Darbois, A.; others. Induction of anergic or regulatory tumor-specific CD4+ T cells in the tumor-draining lymph node. Nature Communications 2018, 9, 1–7.

43. Liu, V.C.; Wong, L.Y.; Jang, T.; Shah, A.H.; Park, I.; Yang, X.; Zhang, Q.; Lonning, S.; Teicher, B.A.; Lee, C. Tumor evasion of the immune system by converting CD4+ CD25− T cells into CD4+ CD25+ T regulatory cells: role of tumor-derived TGF-*β*. The Journal of Immunology 2007, 178, 2883–2892.

44. Vignali, D.A.; Collison, L.W.; Workman, C.J. How regulatory T cells work. Nature Reviews Immunology 2008, 8, 523–532.

45. Batlle, E.; Massagué, J. Transforming Growth Factor-*β* signaling in immunity and cancer. Immunity 2019, 50, 924–940.

46. Wan, Y.Y.; Flavell, R.A. Identifying Foxp3-expressing suppressor T cells with a bicistronic reporter. Proceedings of the National Academy of Sciences 2005, 102, 5126–5131.

47. Wan, Y.Y.; Flavell, R.A. ‘Yin–Yang’ functions of transforming growth factor-*β* and T regulatory cells in immune regulation. Immunological Reviews 2007, 220, 199–213.

48. McKarns, S.C.; Schwartz, R.H. Distinct effects of TGF-*β*1 on CD4+ and CD8+ T cell survival, division, and IL-2 production: a role for T cell intrinsic Smad3. The Journal of Immunology 2005, 174, 2071–2083.

49. Larmonier, N.; Marron, M.; Zeng, Y.; Cantrell, J.; Romanoski, A.; Sepassi, M.; Thompson, S.; Chen, X.; Andreansky, S.; Katsanis, E. Tumor-derived CD4+ CD25+ regulatory T cell suppression of dendritic cell function involves TGF-*β* and IL-10. Cancer Immunology, Immunotherapy 2007, 56, 48–59.

50. Ren, X.; Ye, F.; Jiang, Z.; Chu, Y.; Xiong, S.; Wang, Y. Involvement of cellular death in TRAIL/DR5-dependent suppression induced by CD4+ CD25+ regulatory T cells. Cell Death and Differentiation 2007, 14, 2076–2084.

51. Zhang, K.X.; Firus, J.; Prieur, B.; Jia, W.; Rennie, P.S. To die or to survive, a fatal question for the destiny of prostate cancer cells after androgen deprivation therapy. Cancers 2011, 3, 1498–1512.

52. Ludewig, B.; Krebs, P.; Junt, T.; Metters, H.; Ford, N.J.; Anderson, R.M.; Bocharov, G. Determining control parameters for dendritic cell-cytotoxic T lymphocyte interaction. European Journal of Immunology 2004, 34, 2407–2418.

53. Bladou, F.; Vessella, R.L.; Buhler, K.R.; Ellis, W.J.; True, L.D.; Lange, P.H. Cell proliferation and apoptosis during prostatic tumor xenograft involution and regrowth after castration. International Journal of Cancer 1996, 67, 785–790.

54. Søgaard, C.K.; Moestue, S.A.; Rye, M.B.; Kim, J.; Nepal, A.; Liabakk, N.B.; Bachke, S.; Bathen, T.F.; Otterlei, M.; Hill, D.K. APIM-peptide targeting PCNA improves the efficacy of docetaxel treatment in the TRAMP mouse model of prostate cancer. Oncotarget 2018, 9, 11752–11766.

55. Eisenberg, M.C.; Jain, H.V. A confidence building exercise in data and identifiability: Modeling cancer chemotherapy as a case study. Journal of theoretical biology 2017, 431, 63–78.

56. Rubin, D.B. Using the SIR algorithm to simulate posterior distributions. Bayesian Statistics 1988, 3, 395–402.

57. Dosne, A.G.; Bergstrand, M.; Harling, K.; Karlsson, M.O. Improving the estimation of parameter uncertainty distributions in nonlinear mixed effects models using sampling importance resampling. Journal of Pharmacokinetics and Pharmacodynamics 2016, 43, 583–596.

58. Raftery, A.E.; Bao, L. Estimating and projecting trends in HIV/AIDS generalized epidemics using incremental mixture importance sampling. Biometrics 2010, 66, 1162–1173.

59. Vanlier, J.; Tiemann, C.A.; Hilbers, P.A.; van Riel, N.A. A Bayesian approach to targeted experiment design. Bioinformatics 2012, 28, 1136–1142.

60. Sobol, I.; Levitan, Y.L. A pseudo-random number generator for personal computers. Computers & Mathematics with Applications 1999, 37, 33–40.

61. Saltelli, A.; Annoni, P.; Azzini, I.; Campolongo, F.; Ratto, M.; Tarantola, S. Variance based sensitivity analysis of model output. Design and estimator for the total sensitivity index. Computer Physics Communications 2010, 181, 259–270.

62. Holland, J.H. Adaptation in Natural and Artificial Systems: An Introductory Analysis with Applications to Biology, Control and Artificial Intelligence; MIT Press: Cambridge, MA, USA, 1992.

63. Wei, Y.; Kehm, R.D.; Goldberg, M.; Terry, M.B. Applications for quantile regression in epidemiology. Current Epidemiology Reports 2019, 6, 191–199.

64. Drake, C.G. Update on prostate cancer vaccines. Cancer Journal 2011, 17, 294–299.

65. Anguille, S.; Smits, E.L.; Lion, E.; van Tendeloo, V.F.; Berneman, Z.N. Clinical use of dendritic cells for cancer therapy. The Lancet Oncology 2014, 15, e257–e267.

66. Gatenby, R.A.; Maini, P.K. Mathematical oncology: cancer summed up. Nature 2003, 421, 321–321.

67. Kelley, R.K.; Gane, E.; Assenat, E.; Siebler, J.; Galle, P.R.; Merle, P.; Hourmand, I.O.; Cleverly, A.; Zhao, Y.; Gueorguieva, I.; Lahn, M.; Faivre, S.; Benhadji, K.A.; Giannelli, G. A phase 2 study of galunisertib (TGF-*β*1 receptor type I inhibitor) and sorafenib in patients with advanced hepatocellular carcinoma. Clinical and Translational Gastroenterology 2019, 10.

68. Manogue, C.; Cotogno, P.; Ledet, E.; Lewis, B.; Wyatt, A.W.; Sartor, O. Biomarkers for programmed death-1 inhibition in prostate cancer. The Oncologist 2019, 24, 444–448.

