## Supplementary Information for "Standing Variations Modeling Captures Inter-Individual Heterogeneity in a Deterministic Model of Prostate Cancer Response to Combination Therapy"

### Supplementary Materials

#### S1 Model Variables and Equations

Supplementary Table S1.1: **Model variables and their dimensions.**

| Species | Name | Units |
|---|---|---|
| $N$ | Androgen-sensitive prostate cancer cells | millions of cells |
| $M$ | Castration-resistant prostate cancer cells | millions of cells |
| $D_a$ | Apoptotic cancer cells | millions of cells |
| $D_n$ | Necrotic cancer cells | millions of cells |
| $A_i$ | Immature dendritic cells in the tumor | millions of cells |
| $A_a$ | Activated antigen presenting cells in the tumor | millions of cells |
| $A_m$ | Mature antigen presenting cells in the tumor | millions of cells |
| $T4_a$ | Activated CD4$^+$ T helper cells in the tumor | millions of cells |
| $T8_a$ | Cytotoxic CD8$^+$ T cells in the tumor | millions of cells |
| $Tr_a$ | Activated regulatory T cells in the tumor | millions of cells |
| $C_{\text{TGF}}$ | Amount of TGF-$\beta$ in the tumor | ng |
| $C_{\text{IL2}}$ | Amount of IL-2 in the tumor | ng |
| $[C_{\text{TGF}}]$ | Concentration of TGF-$\beta$ in the tumor | ng/ml |
| $[C_{\text{IL2}}]$ | Concentration of IL-2 in the tumor | ng/ml |
| $v_{tum}$ | Tumor volume | mm$^3$ |
| $r_{tum}$ | Tumor radius | mm |
| $F$ | Find-me chemokine concentration in circulation | arbitrary units |
| $A_m^c$ | Mature antigen presenting cells in circulation post-vaccine injection | millions of cells |
| $A_m^l$ | Mature antigen presenting cells in lymphoid compartment | millions of cells |
| $T4_n^l$ | Naïve CD4$^+$ T helper cells in lymphoid compartment | millions of cells |
| $T4_a^l$ | Activated CD4$^+$ T helper cells in lymphoid compartment | millions of cells |
| $T8_n^l$ | Naïve CD8$^+$ T cells in lymphoid compartment | millions of cells |
| $T8_a^l$ | Activated CD8$^+$ T cells in lymphoid compartment | millions of cells |
| $Tr_r^l$ | Resting regulatory T cells lymphoid compartment | millions of cells |
| $Tr_a^l$ | Activated regulatory T cells lymphoid compartment | millions of cells |
| $C_{\text{TGF}}^l$ | Amount of TGF-$\beta$ in lymphoid compartment | ng |
| $C_{\text{IL2}}^l$ | Amount of IL-2 in lymphoid compartment | ng |
| $[C_{\text{TGF}}^l]$ | Concentration of TGF-$\beta$ in lymphoid compartment | ng |
| $[C_{\text{IL2}}^l]$ | Concentration of IL-2 in lymphoid compartment | ng |

##### S1.1 Model equations

**Tumor compartment equations**

**Androgen sensitive cancer cells – $N$:** Prior to treatment ($\epsilon_N = 1$), $N$ cells proliferate at a rate $\alpha_N$ and undergo apoptotic and necrotic cell death at rates $\delta_N^a$ and $\delta_N^n$, respectively. Under ADT, $N$ cells cease

Supplementary Table S1.2: **Abbreviations.**

| Abbreviation | Name |
|---|---|
| ADT | Androgen deprivation therapy |
| APC | Antigen presenting cell |
| CTL | Cytotoxic $CD8^+$ T lymphocyte |
| DC | Dendritic cell |
| PCa . | Prostate cancer |
| Th cell | $CD4^+$ T helper cell |
| Treg | $CD4^+$ $CD25^+$ $Foxp3^+$ regulatory T cell |

to proliferate ($\epsilon_N = 0$) and undergo additional ADT-dependent cell death at a rate $\delta_N^{\text{adt}}$. Further, these cells also undergo activated CTL-mediated death that occurs at maximum rate $\delta_I$, but is down-regulated in the presence of TGF-$\beta$ [1]. The functional form of CTL-mediated tumor cell death rate is similar to that used by [2]. To account for fact that the speed of CTL infiltration deceases as tumor volume increases [3], the maximum rate of CTL-induced apoptosis is taken to be a decreasing function of $r_{tum}$, the radius of the tumor. In the last term of equation (S1.1), $K_r$ and $K_{\text{TGF}}^I$ are half-saturation constants, and $K_I$ is the CTL to target cell ratio at which rate of cell kill is half its maximum value.

$$\frac{dN}{dt} = \underbrace{\epsilon_N \alpha_N N}_{\substack{\text{ADT-}\\\text{dependent}\\\text{proliferation}}} - \underbrace{\delta_N^a N}_{\substack{\text{apoptotic}\\\text{death}}} - \underbrace{\delta_N^n N}_{\substack{\text{necrotic}\\\text{death}}} - \underbrace{\delta_N^{\text{adt}} N}_{\substack{\text{ADT-}\\\text{induced}\\\text{death}}} - \underbrace{\frac{\frac{\delta_I}{K_r + r_{tum}}}{1 + \frac{[C_{\text{TGF}}]}{K_{\text{TGF}}^I}} \frac{T8_a N}{T8_a + K_I(N + M + A_a + A_m)}}_{\text{CTL- and TGF-}\beta\text{-mediated death}} . \tag{S1.1}$$

**Castration resistant cancer cells –** $M$**:** Prior to treatment ($\epsilon_M = 1$), $M$ cells proliferate at a rate $\alpha_M$ and undergo apoptotic and necrotic cell death at rates $\delta_M^a$ and $\delta_M^n$, respectively. ADT reduces the proliferation rate by a fraction, $0 \le \epsilon_M \le 1$ and may induce limited additional ADT-dependent cell death at a rate $\delta_M^{\text{adt}}$. In equation (S1.2), we expect $\delta_M^{\text{adt}} \ll \delta_N^{\text{adt}}$. As in the case of $N$ cells, $M$ cells also undergo CTL-mediated death, that is down-regulated in the presence of TGF-$\beta$, and is a function of tumor radius. Both $N$ and $M$ cells are assumed to have identical sensitivity to CTLs.

$$\frac{dM}{dt} = \underbrace{\epsilon_M \alpha_M M}_{\substack{\text{ADT-}\\\text{dependent}\\\text{proliferation}}} - \underbrace{\delta_M^a M}_{\substack{\text{apoptotic}\\\text{death}}} - \underbrace{\delta_M^n M}_{\substack{\text{necrotic}\\\text{death}}} - \underbrace{\delta_M^{\text{adt}} M}_{\substack{\text{ADT-}\\\text{induced}\\\text{death}}} - \underbrace{\frac{\frac{\delta_I}{K_r + r_{tum}}}{1 + \frac{[C_{\text{TGF}}]}{K_{\text{TGF}}^I}} \frac{T8_a M}{T8_a + K_I(N + M + A_a + A_m)}}_{\text{CTL- and TGF-}\beta\text{-mediated death}} . \tag{S1.2}$$

**Apoptotic and necrotic cancer cells –** $D_a$ **and** $D_n$**:** When cancer cells die, they move into the apoptotic or necrotic dead cell compartment, depending on the cause of death. Although ADT primarily induces apoptosis in cancer cells, a small amount of necrosis may still occur [4]. Therefore, a fraction $0 < \epsilon_{\text{adt}} < 1$ of the ADT-mediated deaths are assumed to be necrotic, and $0 < (1 - \epsilon_{\text{adt}}) < 1$, apoptotic. CTL-mediated death is assumed to result in cells entering the apoptotic dead cell compartment. Dead cells are primarily cleared by macrophages [5]. For simplicity, we do not explicitly model macrophages; rather, dead cells are assumed to undergo clearance at a rate proportional to their own number, with constant of proportionality, $\delta_D$. In fact, Maree et al. [6] have shown this to be the case when the number of macrophages is constant. Finally, both apoptotic and necrotic dead cells are also phagocytosed by immature DCs [5] at a rate that is an increasing and saturating function of the total number dead cells, and is also proportional to the number of immature DCs, $A_i$ . This phagocytosis term, together with associated parameters, is explained in further detail in section S1.2.

$$\begin{aligned}\frac{dD_a}{dt} &= \underbrace{\delta_N^a\,N + \delta_M^a\,M}_{\substack{\text{apoptotic}\\ \text{death}}} + \underbrace{(1-\epsilon_{\text{adt}})(\delta_N^{\text{adt}}\,N + \delta_M^{\text{adt}}\,M)}_{\text{ADT-mediated apoptosis}} - \underbrace{\lambda_D\,\frac{A_i\,D_a}{K_D v_{tum} + D_a + D_n}}_{\text{uptake by immature DCs}} - \underbrace{\delta_D\,D_a}_{\text{Clearance}} \\ &\quad + \underbrace{\frac{\frac{\delta_I}{K_r + r_{tum}}}{1 + \frac{[C_{\text{TGF}}]}{K_{\text{TGF}}^I}}\,\frac{T8_a\,(N+M)}{T8_a + K_I(N+M+A_a+A_m)}}_{\text{CTL- and TGF-}\beta\text{-mediated death}}, \qquad \text{(S1.3)}\end{aligned}$$

$$\frac{dD_n}{dt} = \underbrace{\delta_N^n\,N + \delta_M^n\,M}_{\substack{\text{necrotic}\\ \text{death}}} + \underbrace{\epsilon_{\text{adt}}(\delta_N^{\text{adt}}\,N + \delta_M^{\text{adt}}\,M)}_{\text{ADT-mediated necrosis}} - \underbrace{\delta_D\,D_n}_{\text{Clearance}} - \underbrace{\lambda_D\,\frac{A_i\,D_n}{K_D v_{tum} + D_a + D_n}}_{\text{uptake by immature DCs}}. \qquad \text{(S1.4)}$$

**Immature DCs – $A_i$:** As a part of innate immunity, immature DCs constantly survey host tissue, and are assumed to localize at the tumor site at a rate that is proportional to the volume of the tumor, $v_{tum}$, with constant of proportionality $s_{A_i}$. This ensures that as the tumor increases in size, the number of DCs per unit volume of the tumor remains constant, assuming no conversion to a mature or an antigen presenting state. The recruitment of DCs to the tumor is up-regulated in the presence of find-me signals, $F$, released by dying tumor cells [5], with constant of proportionality $s'_{A_i}$. DCs also undergo natural death at a rate $\delta_{A_i}$. When DCs phagocytose apoptotic dead tumor cells, they remain untransformed; however, when they phagocytose necrotic dead tumor cells, DCs transform into APCs [7]. Therefore, the rate of DC-to-APC transformation depends on the rate of necrotic cell phagocytosis, while the immature DC is recovered after phagocytosis of apoptotic cells. The last term in equation (S1.5), which describes immature DC activation by necrotic tumor cells, is explained in further detail in section S1.2.

$$\frac{dA_i}{dt} = \underbrace{s_{A_i} v_{tum} + s'_{A_i} F}_{\substack{\text{find-me signal-}\\ \text{mediated DC source}}} - \underbrace{\delta_{A_i}\,A_i}_{\substack{\text{natural}\\ \text{death}}} - \underbrace{\lambda_D\,\frac{A_i\,D_n}{K_D v_{tum} + D_n + D_a}}_{\substack{\text{activation by phagocytosis of}\\ \text{necrotic tumor cells}}}. \qquad \text{(S1.5)}$$

**Activated and mature antigen presenting cells – $A_a$ and $A_m$:** DCs transform into activated DCs or APCs when they phagocytose necrotic dead cells as described above. These APCs undergo further maturation within the tumor, and transform to a phenotype marked by the up-regulation of co-stimulatory molecules such as CD80/CD86 and expression of various cytokines necessary for the activation of effector T cells [8]. The rate of APC maturation is taken to have a maximum value of $\gamma_{A_a}$, and is down-regulated in the presence of Tregs, via direct cell-to-cell contact [9]. Both activated and mature APCs undergo natural death at rates $\delta_{A_a}$ and $\delta_{A_m}$, respectively. Additional cell death follows recognition of cognate antigens on both cell types by activated CTLs, in a process modulated by TGF-$\beta$. Mature APCs migrate to lymphoid organs [10] at an assumed constant per capita rate, $\mu_{A_m}$. In the last term of equation (S1.6), $[Tr_a] = Tr_a/(A_a + T4_a + T8_a)$ is the ratio of Tregs to the other immune cells with which they interact via direct cell-to-cell contact.

$$\frac{dA_a}{dt} = \underbrace{\lambda_D \frac{A_i\, D_n}{K_D v_{tum} + D_n + D_a}}_{\text{Activation of DCs}} - \underbrace{\delta_{A_a} A_a}_{\substack{\text{natural}\\ \text{death}}} - \underbrace{\frac{\frac{\delta_I}{K_r + r_{tum}}}{1 + \frac{[C_{\mathrm{TGF}}]}{K^I_{\mathrm{TGF}}}} \frac{T8_a\, A_a}{T8_a + K_I(N + M + A_a + A_m)}}_{\text{CTL- and TGF-}\beta\text{-mediated death}}$$

$$- \underbrace{\frac{\gamma_{A_a}}{1 + \frac{[Tr_a]}{K^{\gamma}_{Tr}}} A_a}_{\substack{\text{Treg-modulated}\\ \text{maturation}}}, \tag{S1.6}$$

$$\frac{dA_m}{dt} = \underbrace{\frac{\gamma_{A_a}}{1 + \frac{[Tr_a]}{K^{\gamma}_{Tr}}} A_a}_{\substack{\text{Treg-modulated}\\ \text{maturation}}} - \underbrace{\delta_{A_m} A_m}_{\substack{\text{natural}\\ \text{death}}} - \underbrace{\frac{\frac{\delta_I}{K_r + r_{tum}}}{1 + \frac{[C_{\mathrm{TGF}}]}{K^I_{\mathrm{TGF}}}} \frac{T8_a\, A_m}{T8_a + K_I(N + M + A_a + A_m)}}_{\text{CTL- and TGF-}\beta\text{-mediated death}}$$

$$- \underbrace{\mu_{A_m} A_m}_{\substack{\text{migration to}\\ \text{lymphoid organs}}} . \tag{S1.7}$$

**CD4$^+$ helper T cells – $T4_a$:** Th cells are crucial for achieving an effective and regulated immune response. Th cells migrate to the tumor site at a rate $\mu_{T4}$ post-activation in the lymphoid organs [10, 11] and proliferate in the presence of IL-2 [12] at a rate that is taken to be an increasing and saturating function of the concentration of IL-2 in the tumor, with a maximum value of $\alpha_{L4}$, and half-saturation constant $K_L$. This rate of Th cell proliferation decreases due to down-regulation by TGF-$\beta$ [1]. Here, $K^L_{\mathrm{TGF}}$ is a half-saturation constant. Natural death of activated Th cells occurs at a minimum rate of $\delta'_{T4}$ that increases in the presence of activated Tregs in a contact-dependent manner [13]. This rate of Treg-mediated Th cell death is taken to have a maximum value of $\delta''_{T4}$, and is an increasing and saturating function of $[Tr_a]$, the ratio of Tregs to the other immune cells with which they interact, with half-saturation constant $K^{\delta}_{Tr}$. Finally, TGF-$\beta$ has been shown to induce the conversion of normal and activated CD4$^+$ T cells into Tregs [14, 15, 16] within the tumor microenvironment [14]. Therefore, following [2], we include the last term in equation (S1.8) that captures the conversion of activated Th cells to Tregs. This rate of conversion is taken to be an increasing and saturating function of TGF-$\beta$ with a maximum value of $\gamma_{Tr}$ , with half-saturation constant $K^{Tr}_{\mathrm{TGF}}$.

$$\frac{dT4_a}{dt} = \underbrace{\mu_{T4}\, T4^l_a}_{\substack{\text{migration from}\\ \text{lymphoid}\\ \text{organs}}} + \underbrace{\frac{\alpha_{L4}}{1 + \frac{[C_{\mathrm{TGF}}]}{K^L_{\mathrm{TGF}}}} \frac{[C_{\mathrm{IL2}}]}{K_L + [C_{\mathrm{IL2}}]} T4_a}_{\substack{\text{TGF-}\beta\text{- and IL-2 mediated}\\ \text{proliferation}}} - \underbrace{\left(\delta'_{T4} + \delta''_{T4} \frac{[Tr_a]}{K^{\delta}_{Tr} + [Tr_a]}\right) T4_a}_{\text{Treg-mediated cell death}}$$

$$- \underbrace{\gamma_{Tr} \frac{[C_{\mathrm{TGF}}]}{K^{Tr}_{\mathrm{TGF}} + [C_{\mathrm{TGF}}]} T4_a}_{\substack{\text{TGF-}\beta\text{-mediated}\\ \text{conversion to Tregs}}} . \tag{S1.8}$$

**CD8$^+$ cytotoxic T lymphocytes – $T8_a$:** In our model, activated CTLs are terminally differentiated effector T cells with cytotoxic activity. Like Th cells, activated CTLs are recruited to the tumor site at rate $\mu_{T8}$ post-activation in the lymphoid organs [10, 11] and proliferate in the presence of IL-2 [12], at a maximum rate $\alpha_{L8}$ that is down-regulated in the presence of TGF-$\beta$ [1], as in the case of Th cells above. Natural death of activated CTLs occurs at a minimum rate of $\delta'_{T8}$ that increases in the presence of regulatory T cells in a contact-dependent manner [13]. As in equation (S1.8), this rate of Treg-mediated

CTL death is taken to have a maximum value of $\delta''_{T8}$, and is an increasing and saturating function of $[Tr_a]$.

$$\frac{dT8_a}{dt} = \underbrace{\mu_{T8}\, T8_a^l}_{\substack{\text{migration from}\\ \text{lymphoid}\\ \text{organs}}} + \underbrace{\frac{\alpha_{L8}}{1+\frac{[C_{\rm TGF}]}{K_{\rm TGF}^L}}\frac{[C_{\rm IL2}]}{K_L+[C_{\rm IL2}]}T8_a}_{\substack{\text{TGF-}\beta\text{- and IL-2 mediated}\\ \text{proliferation}}} - \underbrace{\left(\delta'_{T8}+\delta''_{T8}\frac{[Tr_a]}{K_{Tr}^{\delta}+[Tr_a]}\right)T8_a}_{\text{Treg-mediated cell death}}. \quad \text{(S1.9)}$$

**CD4$^+$ CD25$^+$ Foxp3$^+$ regulatory T cells – $Tr_a$:** There is evidence that an increase in Treg numbers in cancer patients is a consequence of both, the recruitment of thymus-derived Tregs [17], and the *de novo* generation of peripheral Tregs from CD4$^+$ T cells under TGF-$\beta$ signaling [17, 18]. Here, we do not distinguish between these subtypes. Activated Tregs are recruited to the tumor site at rate $\mu_{Tr}$ post-activation in the lymphoid organs and may also be generated from activated Th cells already present at the tumor site, in the presence of TGF-$\beta$, as described above. The last two terms in equation (S1.10) describe IL-2-dependent Treg proliferation [12] that occurs with a maximum rate of $\alpha_{Lr}$ and activated Treg death, which is taken to occur at a rate $\delta'_{Tr}$.

$$\frac{dTr_a}{dt} = \underbrace{\mu_{Tr}\, Tr_a^l}_{\substack{\text{migration from}\\ \text{lymphoid}\\ \text{organs}}} + \underbrace{\gamma_{Tr}\frac{[C_{\rm TGF}]}{K_{\rm TGF}^{Tr}+[C_{\rm TGF}]}T4_a}_{\substack{\text{TGF-}\beta\text{-mediated}\\ \text{conversion of CD4}^+\text{ Th cells}}} + \underbrace{\alpha_{Lr}\frac{[C_{\rm IL2}]}{K_L+[C_{\rm IL2}]}Tr_a}_{\substack{\text{IL-2 mediated}\\ \text{proliferation}}} - \underbrace{\delta'_{Tr}\, Tr_a}_{\substack{\text{cell}\\ \text{death}}}. \quad \text{(S1.10)}$$

**TGF-$\beta$ chemokine – $C_{TGF}$ and $[C_{TGF}]$:** Activated Tregs, macrophages, and tumor cells all secrete TGF-$\beta$ [19]. However, since we do not explicitly include macrophages in our model, we assume that their number – and hence, macrophage-dependent TGF-$\beta$ production – is proportional to the total number of dead cells. The TGF-$\beta$ secretion rates for all three cell types are $\alpha_T^{Tr}$, $\alpha_T^D$, and $\alpha_T^C$, respectively. Finally, TGF-$\beta$ is assumed to undergo natural decay at constant rate $\delta_T$. The concentration $[C_{TGF}]$ is given by dividing $C_{TGF}$ by the volume of the tumor in ml.

$$\frac{dC_{\rm TGF}}{dt} = \underbrace{\alpha_T^{Tr}\, Tr_a}_{\substack{\text{source from}\\ \text{Treg cells}}} + \underbrace{\alpha_T^D\,(D_a+D_n)}_{\substack{\text{source from}\\ \text{macrophages}}} + \underbrace{\alpha_T^C\,(N+M)}_{\substack{\text{source from}\\ \text{tumor cells}}} - \underbrace{\delta_T\, C_{\rm TGF}}_{\substack{\text{natural}\\ \text{decay}}}, \quad \text{(S1.11)}$$

$$[C_{\rm TGF}] = \frac{C_{\rm TGF}}{v_{tum}\times 10^{-3}}. \quad \text{(S1.12)}$$

**IL-2 chemokine – $C_{\rm IL2}$ and $[C_{\rm IL2}]$:** Activated Th cells secrete the IL-2 cytokine [12] at a maximum rate $\alpha_{T4}$, which is down-regulated in the presence of TGF-$\beta$ [1], with half-saturation constant $K_{\rm TGF}^{T4}$. Further, IL-2 is assumed to undergo natural decay at constant rate $\delta_L$. The concentration $[C_{\rm IL2}]$ is given by dividing $C_{\rm IL2}$ by the volume of the tumor in ml.

$$\frac{dC_{\rm IL2}}{dt} = \underbrace{\frac{\alpha_{T4}}{1+\frac{[C_{\rm TGF}]}{K_{\rm TGF}^{T4}}}T4_a}_{\substack{\text{TGF-}\beta\text{ -mediated}\\ \text{production by Th cells}}} - \underbrace{\delta_L\, C_{\rm IL2}}_{\substack{\text{natural}\\ \text{decay}}}, \qquad [C_{\rm IL2}] = \frac{C_{\rm IL2}}{v_{tum}\times 10^{-3}}. \quad \text{(S1.13)}$$

**Tumor volume and radius – $v_{tum}$ and $r_{tum}$:** The tumor volume is computed by multiplying the volume $v_x$ of each cell type by its number $x$ within the tumor, and adding. To compute its radius, we assume that the tumor is roughly spherical in shape.

$$v_{tum} = \sum_{\text{cell type}} v_x \, x, \qquad r_{tum} = \left( \frac{3\, v_{tum}}{4\pi} \right)^{1/3}. \tag{S1.14}$$

##### Circulation compartment equations

**Find-me chemokines – $F$:** Dying tumor cells release a number of chemotactic factors such as LysoPC and S1P – collectively referred to as find-me signals – that recruit DCs to the tumor site [10]. In our model, both apoptotic and necrotic cells are assumed to produce these signals at the same rate $\alpha_F$. These chemokines are further assumed to extravasate into circulation instantaneously upon release. Find-me chemokines undergo clearance from circulation at constant rate $\delta_F$. Here, $V_d$ is the assumed volume of distribution of these chemokines in circulation.

$$\frac{dF}{dt} = \underbrace{\frac{\alpha_F\,(D_a + D_n)}{V_d}}_{\substack{\text{source from}\\ \text{dead cells}}} - \underbrace{\delta_F\, F}_{\substack{\text{natural}\\ \text{decay}}}. \tag{S1.15}$$

**Mature antigen presenting cells – $A^c_m$:** DC vaccination comprises the injection of mature APCs into circulation, from where they may undergo clearance or accumulate in highly vascularized organs such as the spleen [20].

$$\frac{dA^c_m}{dt} = \underbrace{f(t)}_{\substack{\text{source from}\\ \text{vaccination}}} - \underbrace{\mu_{A^c_m}\, A^c_m}_{\substack{\text{migration to}\\ \text{lymphoid compartment}}} - \underbrace{\lambda_{A^c_m}\, A^c_m}_{\substack{\text{clearance and transfer}\\ \text{to other compartments}}} \tag{S1.16}$$

##### Lymphoid compartment equations

**Mature antigen presenting cells – $A^l_m$:** Mature APCs migrate from the tumor at rate $\mu_{A_m}$ to the lymphoid organs where they can undergo natural death at a rate $\delta_{A_m}$. We assume that there is no further migration of APCs out of lymphoid tissue given that most migrating DCs die after their arrival here [10]. As in equation (S1.7), activated CTLs recognize cognate antigens on mature APCs, and effect cell death that is modulated by TGF-$\beta$.

$$\frac{dA^l_m}{dt} = \underbrace{\mu_{A^c_m}\, A^c_m}_{\substack{\text{migration from}\\ \text{circulation}}} + \underbrace{\mu_{A_m}\, A_m}_{\substack{\text{migration from}\\ \text{tumor}}} - \underbrace{\delta_{A_m}\, A^l_m}_{\substack{\text{natural}\\ \text{death}}} - \underbrace{\frac{\delta'_I}{1 + \frac{[C^l_{\text{TGF}}]}{K^I_{\text{TGF}}}} \frac{T8^l_a\, A^l_m}{T8^l_a + K_I A^l_m}}_{\text{CTL- and TGF-}\beta\text{-mediated death}}. \tag{S1.17}$$

**Naïve CD4$^+$ T helper cells – $T4^l_n$:** Naïve Th cells localize to the lymphoid tissue at an assumed constant rate $s_{T4}$, and undergo natural death at a rate $\delta_{T4}$. They undergo activation by mature APCs at a rate that is assumed to be proportional to the product of the number of Th cells and APCs and has a maximum value of $\lambda_{T4}$. Specifically, when a naïve Th cell comes in contact with a mature APC that presents the correct antigen for its cell surface receptors, the Th cell will become activated [10]. The formulation of this activation term, together with a description of any associated parameters, is provided in section S1.3.

$$\frac{dT4_n^l}{dt} = \underbrace{s_{T4}}_{\substack{\text{source from}\\\text{thymus}}} - \underbrace{\delta_{T4}\,T4_n^l}_{\substack{\text{natural}\\\text{death}}} - \underbrace{\lambda_{T4}\,\frac{A_m^l\,T4_n^l}{K_T v_{spl} + A_m^l + T4_n^l + T8_n^l + Tr_n^l}}_{\text{activation of Th cells}}. \tag{S1.18}$$

**Activated CD4$^+$ T helper cells – $T4_a^l$:** Naïve Th cells are transformed into activated Th cells when they come in contact with APCs as described above. As in the tumor compartment (see equation (S1.8)), activated Th cells in the lymphoid compartment proliferate in the presence of IL-2, undergo Treg-mediated cell death and TGF-$\beta$-mediated conversion to a regulatory phenotype, and will migrate from the lymphoid organs back to the tumor [10, 11] at an assumed constant per capita rate, $\mu_{T4}$. In the third term of equation (S1.19), $[Tr_a^l] = Tr_a^l/(T4_a^l + T8_a^l)$ is the ratio of Tregs to the other immune cells with which they interact via direct cell-to-cell contact.

$$\begin{aligned}\frac{dT4_a^l}{dt} = {} & \underbrace{\lambda_{T4}\,\frac{A_m^l\,T4_n^l}{K_T v_{spl} + A_m^l + T4_n^l + T8_n^l + Tr_n^l}}_{\text{activation of Th cells}} + \underbrace{\frac{\alpha_{L4}}{1 + \frac{[C_{\mathrm{TGF}}^l]}{K_{\mathrm{TGF}}^L}}\,\frac{[C_{\mathrm{IL2}}^l]}{K_L + [C_{\mathrm{IL2}}^l]}\,T4_a^l}_{\substack{\text{TGF-}\beta\text{- and IL-2 mediated}\\\text{proliferation}}} \\ & - \underbrace{\left(\delta'_{T4} + \delta''_{T4}\frac{[Tr_a^l]}{K_{Tr}^{\delta} + [Tr_a^l]}\right) T4_a^l}_{\text{Treg-mediated cell death}} - \underbrace{\gamma_{Tr}\,\frac{[C_{\mathrm{TGF}}^l]}{K_{\mathrm{TGF}}^{Tr} + [C_{\mathrm{TGF}}^l]}\,T4_a^l}_{\substack{\text{TGF-}\beta\text{-mediated}\\\text{conversion to Treg cells}}} - \underbrace{\mu_{T4}\,T4_a^l}_{\substack{\text{migration}\\\text{to tumor}}}.\end{aligned} \tag{S1.19}$$

**Naïve CD8$^+$ cytotoxic T cells – $T8_n^l$:** Naïve CD8$^+$ T cells localize in lymphoid tissue at an assumed constant rate $s_{T8}$, where they undergo natural death at a rate $\delta_{T8}$. Further, when a naïve CD8$^+$ T cell comes in contact with a mature APC that presents the correct antigen for its cell surface receptors, the CD8$^+$ T cell will become an activated CTL [10]. The formulation of this activation term, together with a description of any associated parameters, is further explained in section S1.3.

$$\frac{dT8_n^l}{dt} = \underbrace{s_{T8}}_{\substack{\text{source from}\\\text{thymus}}} - \underbrace{\delta_{T8}\,T8_n^l}_{\substack{\text{natural}\\\text{death}}} - \underbrace{\lambda_{T8}\,\frac{A_m^l\,T8_n^l}{K_T v_{spl} + A_m^l + T4_n^l + T8_n^l + Tr_n^l}}_{\text{activation of CTLs}}. \tag{S1.20}$$

**Activated CD8$^+$ cytotoxic T cells – $T8_a^l$:** Naïve CD8$^+$ T cells are transformed into activated CTLs when they come in contact with APCs. As in the tumor compartment (see equation (S1.9)), activated CTLs in the lymphoid compartment proliferate in the presence of IL-2, undergo Treg-mediated cell death and will migrate from the lymphoid compartment back to the tumor [10, 11] at an assumed constant per capita rate, $\mu_{T8}$.

$$\frac{dT8_a^l}{dt} = \underbrace{\lambda_{T8}\,\frac{A_m^l\,T8_n^l}{K_T v_{spl} + A_m^l + T4_n^l + T8_n^l + Tr_n^l}}_{\text{activation of CTLs}} + \underbrace{\frac{\alpha_{L8}}{1 + \frac{[C_{\mathrm{TGF}}^l]}{K_{\mathrm{TGF}}^L}}\,\frac{[C_{\mathrm{IL2}}^l]}{K_L + [C_{\mathrm{IL2}}^l]}\,T8_a^l}_{\substack{\text{TGF-}\beta\text{- and IL-2 mediated}\\\text{proliferation}}}$$

$$- \underbrace{\left(\delta'_{T8} + \delta''_{T8} \frac{[Tr^l_a]}{K^\delta_{Tr} + [Tr^l_a]}\right) T8^l_a}_{\text{Treg-mediated cell death}} - \underbrace{\mu_{T8}\, T8^l_a}_{\substack{\text{migration}\\ \text{to tumor}}}. \tag{S1.21}$$

**Resting CD4$^+$ CD25$^-$ Foxp3$^-$ regulatory T cells – $Tr^l_r$:** Resting Tregs localize in lymphoid tissue from the thymus at an assumed constant rate $s_{Tr}$, and they undergo natural death at a rate $\delta_{Tr}$. Like naïve CD8$^+$ or CD4$^+$ T cells, resting Tregs undergo activation by mature APCs following T cell receptor (TCR) stimulation [21, 22, 23]. The formulation of this activation term is further explained in section S1.3.

$$\frac{dTr^l_r}{dt} = \underbrace{s_{Tr}}_{\substack{\text{source from}\\ \text{thymus}}} - \underbrace{\delta_{Tr}\, Tr^l_r}_{\substack{\text{natural}\\ \text{death}}} - \underbrace{\lambda_{Tr} \frac{A^l_m\, Tr^l_r}{K_T v_{spl} + A^l_m + T4^l_n + T8^l_n + Tr^l_n}}_{\text{activation of Treg cells}}. \tag{S1.22}$$

**CD4$^+$ CD25$^+$ Foxp3$^+$ regulatory T cells – $Tr^l_a$:** We do not distinguish between activated Tregs derived from the thymus and those generated de novo. Consequently, activated Tregs are generated in our model both, when resting Tregs come in contact with APCs, as well as when activated CD4$^+$ T cells change phenotype under the influence of TGF-$\beta$, as described in equations (S1.19) and (S1.22). As in the tumor compartment (see equation (S1.10)), activated Tregs in the lymphoid compartment proliferate in the presence of IL-2, undergo cell death and will migrate from the lymphoid organs back to the tumor at an assumed constant per capita rate, $\mu_{Tr}$.

$$\begin{aligned} \frac{dTr^l_a}{dt} = {} & \underbrace{\lambda_{Tr} \frac{A^l_m\, Tr^l_r}{K_T v_{spl} + A^l_m + T4^l_n + T8^l_n + Tr^l_n}}_{\text{activation of Treg cells}} + \underbrace{\gamma_{Tr} \frac{[C^l_{\text{TGF}}]}{K^{Tr}_{\text{TGF}} + [C^l_{\text{TGF}}]} T4^l_a}_{\substack{\text{TGF-}\beta\text{-mediated}\\ \text{conversion of CD4}^+\text{ cells}}} \\ & + \underbrace{\alpha_{Lr} \frac{[C^l_{\text{IL2}}]}{K_L + [C^l_{\text{IL2}}]} Tr^l_a}_{\substack{\text{IL-2 mediated}\\ \text{proliferation}}} - \underbrace{\delta'_{Tr}\, Tr^l_a}_{\substack{\text{cell}\\ \text{death}}} - \underbrace{\mu_{Tr}\, Tr^l_a}_{\substack{\text{migration to}\\ \text{lymphoid organs}}}. \end{aligned} \tag{S1.23}$$

**TGF-$\beta$ chemokine – $C^l_{\text{TGF}}$ and $[C^l_{\text{TGF}}]$:** As in equation (S1.11), TGF-$\beta$ is secreted by activated Tregs and it is assumed to undergo natural decay at constant rate $\delta_T$. The concentration $[C^l_{\text{TGF}}]$ is given by dividing $C^l_{\text{TGF}}$ by the volume of the spleen in ml.

$$\frac{dC^l_{\text{TGF}}}{dt} = \underbrace{\alpha^{Tr}_T\, Tr^l_a}_{\substack{\text{source from}\\ \text{Treg cells}}} - \underbrace{\delta_T\, C^l_{\text{TGF}}}_{\substack{\text{natural}\\ \text{decay}}}, \qquad [C^l_{\text{TGF}}] = \frac{C^l_{\text{TGF}}}{v_{spl}}. \tag{S1.24}$$

**IL-2 chemokine – $C^l_{\text{IL2}}$ and $[C^l_{\text{IL2}}]$:** As in equation (S1.13), IL-2 is secreted by activated Th cells in a TGF-$\beta$-dependent manner, and it undergoes natural decay at constant rate $\delta_L$. The concentration $[C^l_{\text{IL2}}]$ is given by dividing $C^l_{\text{IL2}}$ by the volume of the spleen in ml.

$$\frac{dC^l_{\text{IL2}}}{dt} = \underbrace{\frac{\alpha_{T4}}{1 + \frac{[C^l_{\text{TGF}}]}{K^{T4}_{\text{TGF}}}} T4^l_a}_{\substack{\text{TGF-}\beta\text{ -mediated}\\ \text{production by Th cells}}} - \underbrace{\delta_L\, C^l_{\text{IL2}}}_{\substack{\text{natural}\\ \text{decay}}}, \qquad [C^l_{\text{IL2}}] = \frac{C^l_{\text{IL2}}}{v_{spl}}. \tag{S1.25}$$

#### S1.2 Dead cell phagocytosis by dendritic cells

Once DCs arrive at the site of cell death, they must distinguish alive from dying cells. This process is mediated via 'eat-me' signals expressed by dying cells, the most well-characterized one being phosphatidylserine (PtdSer). PtdSer is recognized by a number of membrane adhesion receptors on DCs, such as Tim4, which result in the tethering of the phagocyte to the dead cell. Binding and activation of additional engulfment receptors subsequently leads to corpse engulfment, internalization and digestion [5]. Mathematical models of dead cell or debris removal by phagocytes – specifically, macrophages – have been previously proposed [6, 24]. These models assume that phagocytosis follows mass action kinetics, with irreversible binding and uptake of dead cells or debris by the macrophages, and allow for the possibility of a single macrophage engulfing multiple dead cells.

In our formulation, the process of dead cell uptake by DCs, and their subsequent activation is approximated as follows. Since phagocytes can only engulf dead cells once appropriate cell surface receptors on both cells have formed complexes, we assume that immature DCs bind reversibly to dead cells to form immature DC-dead cell complexes. These complexes then lead to the engulfment and digestion of the dead cell, and the DC is recovered as an immature cell, if an apoptotic cell was ingested, or as an activated APC if a necrotic dead cell was ingested [7]. That is:

$$D_i \; + \; D_a \; \underset{k_d^{\text{ph}}}{\overset{k_a^{\text{ph}}}{\rightleftharpoons}} \; C_{ia} \; \xrightarrow{\lambda_D} \; D_i,$$

$$D_i \; + \; D_n \; \underset{k_d^{\text{ph}}}{\overset{k_a^{\text{ph}}}{\rightleftharpoons}} \; C_{in} \; \xrightarrow{\lambda_D} \; A_a.$$

Here: $D_i$ and $A_a$ are the numbers of uncomplexed immature DCs and activated APCs, respectively; $D_a$ and $D_n$ are the numbers of apoptotic and necrotic dead cells, respectively; and $C_{ia}$ and $C_{in}$ are the numbers of dendritic cell-apoptotic cell and dendritic cell-necrotic cell complexes, respectively. The constants $k_a^{\text{ph}}$ and $k_d^{\text{ph}}$ represent the rates of association and dissociation, respectively, of immature DCs and dead cells, while $\lambda_D$ represents the rate of dead cell ingestion and digestion once a dead cell has tethered to a DC. In the interest of minimizing the dimensionality of parameter space, phagocytes are assumed to engulf and digest apoptotic and necrotic cells at the same rates. For simplicity, we allow DCs to engulf and process only one dead cell at a time. Finally, assuming mass action kinetics as in [6], the above chemical equations can be translated into the following differential equations:

$$\frac{d[D_i]}{dt} = -\,k_a^{\text{ph}}\,[D_i]\,([D_a]+[D_n]) \,+\, k_d^{\text{ph}}\,([C_{ia}]+[C_{in}]) \,+\, \lambda_D\,[C_{ia}], \tag{S1.26}$$

$$\frac{d[A_a]}{dt} = \lambda_D\,[C_{in}], \tag{S1.27}$$

$$\frac{d[D_a]}{dt} = -\,k_a^{\text{ph}}\,[D_i]\,[D_a] \,+\, k_d^{\text{ph}}\,[C_{ia}], \tag{S1.28}$$

$$\frac{d[D_n]}{dt} = -\,k_a^{\text{ph}}\,[D_i]\,[D_n] \,+\, k_d^{\text{ph}}\,[C_{in}], \tag{S1.29}$$

$$\frac{d[C_{ia}]}{dt} = k_a^{\text{ph}}\,[D_i]\,[D_a] \,-\, (k_d^{\text{ph}}+\lambda_D)\,[C_{ia}], \tag{S1.30}$$

$$\frac{d[C_{in}]}{dt} = k_a^{\text{ph}}\,[D_i]\,[D_n] \,-\, (k_d^{\text{ph}}+\lambda_D)\,[C_{in}], \tag{S1.31}$$

where square brackets denote concentration defined as species per unit tumor volume. We now derive a model for dead cell phagocytosis by DCs and their subsequent transformation into APCs. Following Borghans et al. [25], we change variables from free immature DC density ($[D_i]$) to total (free and complexed) immature DC density defined as:

$$[A_i] \;=\; [D_i] \,+\, [C_{ia}] \,+\, [C_{in}]. \tag{S1.32}$$

With this change of variables, the system of equations (S1.26)-(S1.31) transforms to:

$$\frac{d[A_i]}{dt} = -\lambda_D\,[C_{in}], \tag{S1.33}$$
$$\frac{d[A_a]}{dt} = \lambda_D\,[C_{in}], \tag{S1.34}$$
$$\frac{d[D_a]}{dt} = -k_a^{\text{ph}}\,([A_i]-[C_{ia}]-[C_{in}])\,[D_a] + k_d^{\text{ph}}\,[C_{ia}], \tag{S1.35}$$
$$\frac{d[D_n]}{dt} = -k_a^{\text{ph}}\,([A_i]-[C_{ia}]-[C_{in}])\,[D_n] + k_d^{\text{ph}}\,[C_{in}], \tag{S1.36}$$
$$\frac{d[C_{ia}]}{dt} = k_a^{\text{ph}}\,([A_i]-[C_{ia}]-[C_{in}])\,[D_a] - (k_d^{\text{ph}}+\lambda_D)\,[C_{ia}], \tag{S1.37}$$
$$\frac{d[C_{in}]}{dt} = k_a^{\text{ph}}\,([A_i]-[C_{ia}]-[C_{in}])\,[D_n] - (k_d^{\text{ph}}+\lambda_D)\,[C_{in}]. \tag{S1.38}$$

Given that we are interested on a timescale of prostate xenograft growth in mice that occurs over several weeks, and the fact that DCs typically phagocytose dead cells within 3 hours [7], we assume quasi-steady state kinetics and take the rates of complex formation in the above equations to zero. Then,

$$[C_{ia}] = \frac{[A_i]\,[D]_a}{K_D+[D_a]+[D_n]} \qquad \text{and} \qquad [C_{in}] = \frac{[A_i]\,[D_n]}{K_D+[D_a]+[D_n]}, \tag{S1.39}$$

where $K_D = (k_d^{\text{ph}}+\lambda_D)/k_a^{\text{ph}}$. Substituting these in the equations (S1.35)-(S1.36), and using the quasi-steady state assumption on $[C_{ia}]$ and $[C_{in}]$ dynamics, we deduce that the rates of apoptotic and necrotic cell phagocytosis by DCs are:

$$\frac{d[D_a]}{dt} = -\lambda_D\,\frac{[A_i]\,[D_a]}{K_D+[D_a]+[D_n]}, \qquad \text{and} \qquad \frac{d[D_n]}{dt} = -\lambda_D\,\frac{[A_i]\,[D_n]}{K_D+[D_a]+[D_n]}. \tag{S1.40}$$

Likewise, substituting equations (S1.39) in the equations (S1.33)-(S1.34) allows us to deduce the rate at which DCs transform into antigen presenting cells as follows:

$$\frac{d[A_i]}{dt} = -\lambda_D\,\frac{[A_i]\,[D_n]}{K_D+[D_a]+[D_n]}, \qquad \text{and} \qquad \frac{d[A_m]}{dt} = \lambda_D\,\frac{[A_i]\,[D_n]}{K_D+[D_a]+[D_n]}. \tag{S1.41}$$

Finally, we convert from concentration of cellular species back to total cell numbers by observing that [variable] = variable/$v_{tum}$, where $v_{tum}$ is tumor volume, and making the simplifying assumption that $v_{tum} \approx$ constant on the time-scale of tumor cell phagocytosis. Then:

$$\frac{dD_a}{dt} = -\lambda_D\,\frac{A_i\,D_a}{K_D v_{tum}+D_a+D_n}, \qquad \frac{dD_n}{dt} = -\lambda_D\,\frac{A_i\,D_n}{K_D v_{tum}+D_a+D_n},$$
$$\frac{dA_i}{dt} = -\lambda_D\,\frac{A_i\,D_n}{K_D v_{tum}+D_a+D_n}, \qquad \frac{dA_m}{dt} = \lambda_D\,\frac{A_i\,D_n}{K_D v_{tum}+D_a+D_n}. \tag{S1.42}$$

#### S1.3 T cell activation by antigen presenting cells

The process of T cell activation by activated DCs, or APCs, is highly complex, involving several cytokines, cell surface receptors, co-factors and various subsets of the T cell population such as CTLs, Th cells and Tregs; however, incorporating extensive biological detail in the model would necessitate the addition of several unknown parameters. We therefore focus on those aspects of the APC-mediated T cell-activation pathway most relevant to understanding the response of prostate cancer to vaccination therapy. Briefly, APCs travel from the tumor tissue to lymphoid organs such as the spleen and lymph nodes where they activate naïve, CD4- and CD8-expressing T cells, transforming them into Th cells and

CTLs, respectively. T cell antigen receptors on T cells recognize fragments of tumor-derived antigens bound to MHC-I complexes on APCs, resulting in a transient APC-naïve T cell complex. Co-stimulatory signals, such as ICAM-1, CD58 and B7, expressed by the same APC bind to various molecules such as LFA-1, CD2 and CD58 on the T cell, enhancing APC-T cell adhesion and subsequent activation of the T lymphocyte [10].

Several mathematical models of T cell activation by APCs have been proposed (see [26] for a recent review of these). Here, we follow the approach of De Boer and Perelson [27]. In their model, APCs bind reversibly to naïve T cells, resulting in a T cell-APC complex that subsequently yields an activated T cell and the APC is recovered. The activated T cell subsequently divides into two daughter cells that are either naïve or 'experienced' – but not activated. The model also allows for distinction between various T cell sub-populations. We modify this approach slightly as follows. For simplicity, we only consider a single naïve T cell population for each cell type, that is, $CD4^+$, $CD8^+$ and resting Tregs. Further, we model activated T cell proliferation as a separated event since, biologically, T cell proliferation is dependent on interleukin-2 (IL-2), primarily expressed by APC-activated $CD4^+$ T helper cells [12]. That is, the following biochemical equations are assumed to represent APC-mediated Th and CTL activation:

$$
\begin{aligned}
T4^l_{n,f} \;+\; A^l_{m,f} \;&\underset{k^{\text{ac}}_{d4}}{\overset{k^{\text{ac}}_{a4}}{\rightleftharpoons}}\; C_{4a} \;\xrightarrow{\lambda_{T4}}\; T4^l_a \;+\; A^l_{m,f},\\
T8^l_{n,f} \;+\; A^l_{m,f} \;&\underset{k^{\text{ac}}_{d8}}{\overset{k^{\text{ac}}_{a8}}{\rightleftharpoons}}\; C_{8a} \;\xrightarrow{\lambda_{T8}}\; T8^l_a \;+\; A^l_{m,f},\\
Tr^l_{r,f} \;+\; A^l_{m,f} \;&\underset{k^{\text{ac}}_{dr}}{\overset{k^{\text{ac}}_{ar}}{\rightleftharpoons}}\; C_{ra} \;\xrightarrow{\lambda_{Tr}}\; Tr^l_a \;+\; A^l_{m,f}.
\end{aligned}
$$

Here: $T4^l_{n,f}$ and $T4^l_a$ are the numbers of naïve, uncomplexed Th cells, and activated Th cells, respectively; $T8^l_{n,f}$ and $T8^l_a$ are the numbers of naïve, uncomplexed CTLs, and activated CTLs, respectively; $Tr^l_{r,f}$ and $Tr^l_a$ are the numbers of resting, uncomplexed Tregs, and activated Tregs, respectively; $A^l_{m,f}$ is the number of free, antigen-presenting APCs; and $C_{4a}$, $C_{8a}$ and $C_{ra}$ are the numbers of naïve Th-APC cell complexes, naïve CTL-APC cell complexes and resting Treg-APC cell complexes, respectively. The superscript $l$ denotes the fact that these reactions are occurring in lymphoid tissue. The constants $k^{\text{ac}}_{a*}$ and $k^{\text{ac}}_{d*}$ represent the rates of association and dissociation, respectively, of naïve or resting T cells with APCs while $\lambda_{T*}$ represents the rate of APC unbinding from the complex, resulting in an activated Th cell, CTL or Treg ($* = 4$, 8 or $r$). The above reaction scheme is essentially an enzyme-catalyzed reaction with the APCs playing the part of the enzyme, and the naïve and activated T cells playing the part of the substrate and product, respectively. Such reactions have been studied extensively (see [28] for a recent review). Here, we follow the approach of Borghans et al. [25] wherein mass action kinetics are assumed, and a total quasi steady state approximation made that is valid for a broad range of parameter values including high and low enzyme – or, in our case, APC – concentrations.

Briefly, the variables representing free, naïve Ths ($T4^l_{n,f}$), free, naïve CTLs ($T8^l_{n,f}$), resting, free Tregs ($Tr^l_{r,f}$) and free APCs ($A^l_{m,f}$) are replaced by three new variables $T4^l_n$, $T8^l_n$, $Tr^l_r$ and $APC^l$ (defined below), representing the total numbers of naïve Ths, naïve CTLs, resting Tregs and APCs, respectively.

$$
\begin{aligned}
[T4^l_n] \;&=\; [T4^l_{n,f}] + [C_{4a}], \qquad & [T8^l_n] \;&=\; [T8^l_{n,f}] + [C_{8a}],\\
\left[Tr^l_r\right] \;&=\; [Tr^l_{r,f}] + [C_{ra}], \qquad & [A^l_m] \;&=\; [A^l_{m,f}] + [C_{4a}] + [C_{8a}] + [C_{ra}],
\end{aligned} \tag{S1.43}
$$

where square brackets denote concentration defined as species per unit spleen volume. Further, the rates of formation of $[C_{4a}]$, $[C_{8a}]$ and $[C_{ra}]$ are assumed to be at a quasi steady state, yielding the following equations:

$$
0 \;=\; k^{\text{ac}}_{a4}\left([T4^l_n] - [C_{4a}]\right)\left([A^l_m] - [C_{4a}] - [C_{8a}] - [C_{ra}]\right) - (k^{\text{ac}}_{d4} + \lambda_{T4})\,[C_{4a}], \tag{S1.44}
$$

$$0 = k^{\rm ac}_{a8}\Big([T8^l_n] - [C_{8a}]\Big)\Big([A^l_m] - [C_{4a}] - [C_{8a}] - [C_{ra}]\Big) - (k^{\rm ac}_{d8} + \lambda_{T8})\,[C_{8a}], \quad \text{(S1.45)}$$

$$0 = k^{\rm ac}_{ar}\Big([Tr^l_r] - [C_{ra}]\Big)\Big([A^l_m] - [C_{4a}] - [C_{8a}] - [C_{ra}]\Big) - (k^{\rm ac}_{dr} + \lambda_{Tr})\,[C_{ra}], \quad \text{(S1.46)}$$

$$\frac{d[T4^l_n]}{dt} = -\,\lambda_{T4}\,[C_{4a}], \quad \text{(S1.47)}$$

$$\frac{d[T4^l_a]}{dt} = \lambda_{T4}\,[C_{4a}], \quad \text{(S1.48)}$$

$$\frac{d[T8^l_n]}{dt} = -\,\lambda_{T8}\,[C_{8a}], \quad \text{(S1.49)}$$

$$\frac{d[T8^l_a]}{dt} = \lambda_{T8}\,[C_{8a}], \quad \text{(S1.50)}$$

$$\frac{d[Tr^l_r]}{dt} = -\,\lambda_{Tr}\,[C_{ra}], \quad \text{(S1.51)}$$

$$\frac{d[Tr^l_a]}{dt} = \lambda_{Tr}\,[C_{ra}], \quad \text{(S1.52)}$$

and where $[A^l_m]$ is a conserved quantity. Applying an argument similar to that in [25], the following approximate solutions to equations (S1.44)-(S1.46) may be derived:

$$[C_{4a}] = \frac{[A^l_m]\,[T4^l_n]\,([A^l_m] + K_{T8})\,([A^l_m] + K_{Tr})}{Denom}, \quad \text{(S1.53)}$$

$$[C_{8a}] = \frac{[A^l_m]\,[T8^l_n]\,([A^l_m] + K_{T4})\,([A^l_m] + K_{Tr})}{Denom}, \quad \text{(S1.54)}$$

$$[C_{ra}] = \frac{[A^l_m]\,[Tr^l_r]\,([A^l_m] + K_{T4})\,([A^l_m] + K_{T8})}{Denom}, \quad \text{(S1.55)}$$

where:

$$Denom = K_{T4}K_{T8}K_{Tr} + \sum_{i\neq j\neq k}[Ti^l_*]K_{Tj}K_{Tk} + [A^l_m]\Big(\sum_{i\neq j}K_{Ti}K_{Tj} + \sum_{i\neq j\neq k}[Ti^l_*]\big(K_{Tj} + K_{Tk}\big)\Big)$$
$$+\,[A^l_m]^2\Big(\sum_i([Ti^l_*] + K_{Ti})\Big) + [A^l_m]^3, \qquad i,j,k \in \{4,8,r\},\ * = n \text{ or } r, \quad \text{(S1.56)}$$

and where $K_{T4} = (k^{\rm ac}_{d4} + \lambda_{T4})/k^{\rm ac}_{a4}$, $K_{T8} = (k^{\rm ac}_{d8} + \lambda_{T8})/k^{\rm ac}_{a8}$ and $K_{Tr} = (k^{\rm ac}_{dr} + \lambda_{Tr})/k^{\rm ac}_{ar}$. We remark that the above expressions are needlessly complex. We therefore make the simplifying assumption that $K_{T4} \approx K_{T8} \approx K_{Tr}\,(= K_T$, say). That is, T cell receptors on naïve Ths, CTLs and resting Tregs are equally sensitive to binding MHC-I complexes on APCs. Under this assumption, the above expressions simplify to yield:

$$[C_{4a}] \approx \frac{[A^l_m]\,[T4^l_n]}{K_T + [A^l_m] + [T4^l_n] + [T8^l_n] + [Tr^l_r]}, \qquad [C_{8a}] \approx \frac{[A^l_m]\,[T8^l_n]}{K_T + [A^l_m] + [T4^l_n] + [T8^l_n] + [Tr^l_r]},$$
$$[C_{ra}] \approx \frac{[A^l_m]\,[Tr^l_r]}{K_T + [A^l_m] + [T4^l_n] + [T8^l_n] + [Tr^l_r]}. \quad \text{(S1.57)}$$

Finally, we convert from concentration of cellular species back to total cell numbers by observing that [variable] = variable/$v_{spl}$, where $v_{spl}$ is spleen volume, which is constant over time. Then, the equations governing the rates of change of T cell numbers are:

$$\frac{dTi^l_*}{dt} = -\,\lambda_{Ti}\,\frac{A^l_m\,Ti^l_*}{K_T v_{spl} + A^l_m + T4^l_n + T8^l_n + Tr^l_r}, \quad \text{(S1.58)}$$

$$\frac{dTi^l_a}{dt} = \lambda_{Ti}\,\frac{A^l_m\,Ti^l_*}{K_T v_{spl} + A^l_m + T4^l_n + T8^l_n + Tr^l_r}, \quad \text{(S1.59)}$$

where $i = 4, 8$ and $* = n$ or $i = r$ and $* = r$.

### S2 Parameter Estimation

Supplementary Table S2.1: **Parameter values taken from the literature.**

| **Parameter** | **Meaning** | **Value** | **Units** | Source |
|---|---|---|---|---|
| $v_C$ | Volume of $10^6$ tumor cells | 1.2542 | $mm^3$ | [29] |
| $v_{D_a}$ | Volume of $10^6$ apoptotic tumor cells | $0.7\,v_C$ | $mm^3$ | [30] |
| $v_{D_n}$ | Volume of $10^6$ necrotic tumor cells | $1.3\,v_C$ | $mm^3$ | [31, 32] |
| $v_A$ | Volume of $10^6$ DCs or APCs | 0.7 | $mm^3$ | [33] |
| $v_T$ | Volume of $10^6$ T cells | 0.176 | $mm^3$ | [34] |
| $\delta_{A_i}$ | Death rate of immature DCs | 0.3151 | per day | [35] |
| $\delta_{A_a}$ | Death rate of activated APCs | 0.40 | per day | [36] |
| $\delta_{A_m}$ | Death rate of mature APCs | 0.40 | per day | [36] |
| $\delta_{T4}$ | Death rate of naïve Th cells | 0.0115 | per day | [37] |
| $\delta'_{T4}$ | Death rate of activated Th cells | 0.1199 | per day | see text |
| $\delta''_{T4}$ | Treg-mediated death rate of activated Th cells | 0.3421 | per day | see text |
| $\delta_{T8}$ | Death rate of naïve CTLs | 0.0130 | per day | [37] |
| $\delta'_{T8}$ | Death rate of activated CTLs | 0.1199 | per day | [38] |
| $\delta''_{T8}$ | Treg-mediated death rate of activated CTLs | 0.3421 | per day | see text |
| $\delta_{Tr}$ | Death rate of resting Treg cells | 0.0115 | per day | [37, 39] |
| $\delta_I$ | CTL-induced death rate of tumor cells | 24.1920 | per day | [3, 40] |
| $\delta_T$ | TGF-$\beta$ degradation rate | 8.640 | per day | [41] |
| $\delta_L$ | IL-2 degradation rate | 12.50 | per day | [2] |
| $\gamma_{A_a}$ | Active APC maturation rate | 1.70 | per day | [42] |
| $\mu_{A_m}$ | Rate of mature APC transfer from tumor to lymphoid compartment | 1.3333 | per day | [43] |

Where possible, parameter values were taken from the literature. These are listed in Tables S2.1 and S2.2 together with their sources. The rates of naïve T cell localization to the mouse spleen were estimated as follows. A typical healthy mouse spleen weighs between 80 and 120 mg and contains up to $10^8$ splenocytes[1]. In the JAX-FVB-NJ mouse used in the experiments of Shen at el. [44], 6-9% of splenocytes are $CD8^+$ T cells and 18-25% are $CD4^+$ T cells[2]. Also, under healthy homeostasis, 6-10% of $CD4^+$ T cells are Tregs [45]. From this information, healthy steady-state levels of the various naïve T cell populations in the spleen can be estimated. Assuming that, in the absence of a tumor there are no activated T cells, the rate of $i$-type T cell localization to the spleen ($s_i$) can be calculated as $s_i = i_\infty * \delta_i$, where $i_\infty$ is the steady state number of $i$-type T cells in the spleen, and $\delta_i$, their natural death rate. Further, the death rates of activated and mature APCs were assumed to be identical, that is, $\delta_{A_a} = \delta_{A_m}$. The maximum value of Treg-mediated CTL death rate, $\delta''_{T8}$, was obtained as follows. Radunskaya et al. [38] estimate the death rate of tumoral CTLs to be 0.4620 per day and the death rate of newly activated lymphoid CTLs, $\delta'_{T8}$, to be 0.1199 per day. Since Radunskaya et al. do not consider Tregs in their model, we assume that 0.4620 per day is the maximum possible death rate of activated CTLs, that is, $\delta'_{T8} + \delta''_{T8} = 0.4620$. From this, $\delta''_{T8}$ may be estimated. Next, following a similar assumption in [2], the constitutive and Treg-induced death rates of activated Th cells are assumed to be identical to those of

[1] `https://www.miltenyibiotec.com/US-en/resources/macs-handbook/mouse-cells-and-organs/mouse-cell-types/cd4-t-cells-mouse.html`

[2] `http://jackson.jax.org/rs/444-BUH-304/images/physiological_data_001800.pdf`

activated CTLs, that is, $\delta'_{T4} = \delta'_{T8}$ and $\delta''_{T4} = \delta''_{T8}$. Finally, the maximum rate of tumor cell kill by CTLs, $\delta_I$, was estimated as follows. Once CTLs arrive at the tumor, it takes them between 5-6 days to diffuse through the tumor, migrating at a speed of 12.096 mm per day [3]. Further, it takes a CTL on average 6 hours to induce apoptosis in a target cell [40]. From this, $\delta_I$ and $K_r$ may be estimated. Note that this yields an upper bound for $\delta_I$, since the precise value of this parameter will be influenced by factors such as CTL specificity for tumor cells, and other tumor-mediated immune-inhibitory mechanisms not explicitly considered here, such as the expression of programmed death-ligand 1 (PD-L1) on cancer cells.

Supplementary Table S2.2: **Parameter values taken from the literature.**

| **Parameter** | **Meaning** | **Value** | **Units** | Source |
|---|---|---|---|---|
| $K^I_{\text{TGF}}$ | Half-saturation constant for TGF-$\beta$-inhibited tumor cell kill | 3.50 | ng/ml | [2] |
| $K^L_{\text{TGF}}$ | Half-saturation constant for TGF-$\beta$-inhibited T cell proliferation | 2.90 | ng/ml | [2] |
| $K^{Tr}_{\text{TGF}}$ | Half-saturation constant for TGF-$\beta$-mediated Th-to-Treg conversion | 1.70 | ng/ml | [2] |
| $K^{T4}_{\text{TGF}}$ | Half-saturation constant for TGF-$\beta$-inhibited IL-2 production | 0.90 | ng/ml | [2] |
| $K^{\delta}_{Tr}$ | Half-saturation constant for Treg-mediated death of activated T cells | 0.7873 | dimensionless | [13] |
| $K^{\gamma}_{Tr}$ | Half-saturation constant for Treg-mediated inhibition of APC maturation | 0.8517 | dimensionless | [9] |
| $K_L$ | Half-saturation constant for IL-2-mediated active T cell proliferation | 0.3 | ng/ml | [2] |
| $K_r$ | Half-saturation constant for effect of tumor size on CTL-mediated tumor cell death rate | 6.0480 | mm | [3, 40] |
| $\alpha_{L4}$ | Maximum rate of activated Th cell proliferation | 1.47 | per day | [46] |
| $\alpha_{L8}$ | Maximum rate of activated CTL proliferation | 1.6094 | per day | [47] |
| $\alpha_{Lr}$ | Maximum rate of activated Treg proliferation | 2.10 | per day | [2] |
| $\alpha^{Tr}_{T}$ | Treg production rate of TGF-$\beta$ | 3.7544 | ng per million cells per day | [14, 48] |
| $\alpha_{T4}$ | Th cell production rate of IL-2 | 10.14 | ng per million cells per day | [1] |
| $s_{T4}$ | Localization rate of naïve Th cells in spleen | 0.2460 | million cells per day | see text |
| $s_{T8}$ | Localization rate of naïve CTLs in spleen | 0.0975 | million cells per day | see text |
| $s_{Tr}$ | Localization rate of resting Tregs in spleen | 0.0197 | million cells per day | see text |

Next , we describe the estimation of parameter values that were fit to experimental data. These are listed in Table S2.3, and best fits plotted in Figure 2 of the main text. Parameters relating to tumor cell proliferation and death, and dead cell clearance were estimated as follows. In a series of experiments by Shen et al. reported in [44], prostate cancer xenografts comprising the Myc-Cap cell line were established in 8-10 week-old male FVB/NJ mice. Tumor diameters were measured periodically, and ADT started on day 22, when tumor volumes reached approximately 400 mm$^3$. The onset of castration resistance was observed, and defined as when tumor volumes increased to a minimum of 420 mm$^3$ after the initial decline post-ADT initiation. The emergence of castration resistance in the Myc-Cap xenograft model occurs due to an amplification of the androgen receptor gene [49]. Here, we approximate this process by assuming that there is an initial pool of castration-resistant cells already present at the time of ADT initiation. The time-course tumor volume data pre- and post-ADT described above was used to estimate the rates of tumor cell proliferation $\alpha_N$ and $\alpha_M$, the ADT-induced death rate of androgen sensitive cells $\delta_N^{\text{adt}}$, and the rate of dead cell clearance, $\delta_D$. We remark that, for simplicity, androgen deprivation is taken to have no effect on castration resistant cells, that is, in equation (S1.2), $\epsilon_M = 1$ and $\delta_M^{\text{adt}} = 0$. The results of these fits are shown in Figure 2A. Further, Bladou et al. [50] have estimated that roughly 0.66% of cancer cells in prostate cancer xenografts are apoptotic in the absence of any treatment, from which the constitutive or background rates of cancer cell apoptosis, $\delta_N^a$ and $\delta_M^a$ were estimated. Soggard et al. [51] have estimated that equal fractions of untreated prostate cancer xenografts are apoptotic and necrotic. Therefore, we take the constitutive or background rates of cancer cell necrosis apoptosis to be equal, that is, $\delta_N^n = \delta_N^a$ and $\delta_M^n = \delta_M^a$. The results of these fits are shown in Figure 2B.

Supplementary Table S2.3: **Parameter values fit to experimental data.**

| **Parameter** | **Meaning** | **Value** | **Units** |
|---|---|---|---|
| $\alpha_N$ | Rate of $N$ cell proliferation | 0.1473 | per day |
| $\alpha_M$ | Rate of $M$ cell proliferation | 0.1862 | per day |
| $\delta_N^a$ | Rate of $N$ cell apoptosis | 0.0074 | per day |
| $\delta_N^n$ | Rate of $N$ cell necrosis | 0.0074 | per day |
| $\delta_N^{\text{adt}}$ | Rate of ADT-mediated $N$ cell death | 0.0687 | per day |
| $\delta_M^a$ | Rate of $M$ cell apoptosis | 0.0093 | per day |
| $\delta_M^n$ | Rate of $M$ cell necrosis | 0.0093 | per day |
| $\delta_M^{\text{adt}}$ | Rate of ADT-mediated $M$ cell death | 0 | per day |
| $\delta'_{Tr}$ | Rate of activated Treg death | 1.0216 | per day |
| $\delta_D$ | Rate of dead cell clearance | 0.9738 | per day |
| $s_{Ai}$ | Constitutive localization rate of DCs to tumor | $3.620\times10^{-3}$ | millions of cells per gm per day |
| $s'_{Ai}$ | Find-me signal-mediated localization rate of DCs to tumor | $0.110\times10^{-3}$ | DCs per dead cell per day |
| $\lambda_{T4}$ | Maximum rate of CTL activation by APCs | 4.1584 | per day |
| $\lambda_{T8}$ | Maximum rate of Th cell activation by APCs | 0.7908 | per day |
| $\alpha_T^C$ | Production rate of TGF-$\beta$ by cancer cells | $7.038\times10^{-2}$ | ng per million cells per day |
| $\alpha_T^D$ | Production rate of TGF-$\beta$ by macrophages | 4.4545 | ng per million cells per day |
| $\gamma_{Tr}$ | Maximum rate of TGF-$\beta$-mediated Th-to-Treg conversion | 1.5944 | per day |

In the same set of experiments by Shen et al. [44] described above, the immune presence at the site of the xenograft was also recorded. Specifically, the total numbers of DCs (immature, activated and mature), CD4+ T cells (Th cells and Tregs), CD8+ T Cells, and Tregs were counted prior to ADT initiation, 7 days post ADT initiation, and once castration resistance had emerged. This data was used to estimate the following parameters: immature DC localization rates at the tumor site, $s_{A_i}$ and $s'_{A_i}$; activation rates of T cells by mature APCs in the lymphoid compartment, $\lambda_{T4}$, $\lambda_{T8}$ and $\lambda_{Tr}$; the death rate of activated Tregs, $\delta'_{Tr}$; TGF-$\beta$-mediated rate of Th cell conversion to Treg, $\gamma_{Tr}$ ; and the rates of TGF-$\beta$ expression by tumor cells and macrophages, $\alpha_T^C$ and $\alpha_T^D$, respectively. The results of these fits are shown in Figure 2C-E. We remark that in the absence of data with which to estimate the parameters relating to find-me signal production (see equation (S1.15)), we assume that the concentration of $F$ in circulation is at a quasi steady-state, that is:

$$F = \alpha'_F (D_a + D_n), \qquad \text{where} \qquad \alpha'_F = \alpha_F / V_d \, \delta_F. \tag{S2.60}$$

The above expression for $F$ is then substituted in equation (S1.5), and the unknown parameter $\alpha'_F$ absorbed in $s'_{A_i}$, the find-me signal-mediated rate of DC localization to the tumor. That is,

$$\frac{dA_i}{dt} = \underbrace{s_{A_i} v_{tum} + s'_{A_i}(D_a + D_n)}_{\substack{\text{find-me signal-} \\ \text{mediated DC source}}} - \underbrace{\delta_{A_i} A_i}_{\substack{\text{natural} \\ \text{death}}} - \underbrace{\lambda_D \frac{A_i D_n}{K_D v_{tum} + D_n + D_a}}_{\substack{\text{activation by phagocytosis of} \\ \text{necrotic tumor cells}}}. \quad \text{(S2.61)}$$

We list in Table S2.4 values of those parameters, which could not be determined from the literature. In these cases, biologically realistic values were assigned to these parameters. For instance, the rates of activated T cell trafficking to t he tumor, $\mu_{T4}$, $\mu_{T8}$ and $\mu_{Tr}$, are assumed to be equal to the rate of mature APC transfer from the tumor to the lymphoid compartment, $\mu_{A_m}$. A value for the fraction $\epsilon_{\text{adt}}$ of ADT-induced tumor cell death that is necrotic was chosen keeping in mind that ADT primarily induces apoptosis in cancer cells [4]. Further, it is estimated that immature DCs take an average of 3 hours to phagocytose dead cells [7]. Consequently, we assume that the rate of dead cell phagocytosis by DCs, and subsequent DC activation, $\lambda_D$, is no larger than 8 per day. Finally, $\lambda_{Tr}$ – the maximum rate of Treg activation by mature APCs – is taken to be the same as that of CTLs, that is, $\lambda_{T8}$.

Supplementary Table S2.4: **Additional parameter values.**

| **Parameter** | **Meaning** | **Value** | **Units** |
|---|---|---|---|
| $\epsilon_{\text{adt}}$ | Fraction of ADT-induced cell death that is necrotic | 0.05 | dimensionless |
| $\mu_{T4}$ | Rate of activated Th cell migration to tumor | 1.3333 | per day |
| $\mu_{T8}$ | Rate of activated CTL migration to tumor | 1.3333 | per day |
| $\mu_{Tr}$ | Rate of activated Treg migration to tumor | 1.3333 | per day |
| $K_D$ | Half-saturation constant for dead cell phagocytosis by DCs | 0.01 | millions of cells per mm$^3$ |
| $K_T$ | Half-saturation constant for APC-mediated activation of T cells | 20 | millions of cells per mm$^3$ |
| $\lambda_D$ | Maximum rate of dead cell phagocytosis by DCs, and of DC conversion to APC | 6 | per day |
| $\lambda_{Tr}$ | Maximum rate of Treg activation by APCs | 4.1584 | per day |

We next used standard errors reported in the experimental data in [44] to quantify the uncertainty in a subset of the parameters listed in Tables S2.3 and S2.4, using our standing variations approach. A total of 18 parameters were selected for this analysis, with the resultant distributions plotted in Figures S2.2 and S2.3, and their 95% bounds and median values recorded in Table S2.5. Finally, one key parameter in our model is $K_I$, the CTL to target cell ratio at which rate of CTL-induced cell kill is half its maximum value. This determines the effectiveness of CTLs at targeting tumor cells. However, we do not have direct experimental data with which to estimate its value. We therefore assumed that it follows a unimodal Weibull-Normal distribution [52] across the mouse population we are modeling.

Supplementary Table S2.5: **95% confidence bounds on sampled parameters.**

| **Tumor growth and response to ADT** | | |
|---|---|---|
| **Parameter** | **Median value** | **Confidence bounds** |
| $v_{tum}(t = 10)$ | 91.7190 | $[52.13, 133.84]$ |
| $\alpha_N$ | 0.1587 | $[0.0990, 0.2251]$ |
| $\alpha_M$ | 0.1559 | $[0.1277, 0.2088]$ |
| $\delta_N^{\text{adt}}$ | 0.0969 | $[0.0478, 0.1696]$ |
| $\delta_D$ | 2.4038 | $[0.2516, 4.7461]$ |
| $M_0 = M(t = 10)$ | 1.7292 | $[0.2171, 4.9340]$ |
| **Immune presence in tumor** | | |
| **Parameter** | **Median value** | **Confidence bounds** |
| $s_{Ai}$ | $4.7351\times10^{-3}$ | $[2.3983, 7.0021] \times 10^{-3}$ |
| $s'_{Ai}$ | $0.0864\times10^{-3}$ | $[0.0125, 0.2619] \times 10^{-3}$ |
| $\lambda_{T4}$ | 5.3770 | $[2.3597, 7.1985]$ |
| $\lambda_{T8}$ | 1.2050 | $[0.5247, 1.8564]$ |
| $\gamma_{Tr}$ | 1.3521 | $[0.4322, 3.2551]$ |
| $\alpha_T^C$ | $3.7524\times10^{-2}$ | $[0.3795, 11.2970] \times 10^{-2}$ |
| $\alpha_T^D$ | 5.2869 | $[1.0288, 8.2441]$ |
| $\delta'_{Tr}$ | 1.2844 | $[0.4334, 1.9986]$ |
| $\epsilon_{\text{adt}}$ | 0.1722 | $[0.0176, 0.3889]$ |
| $K_D$ | $1.0922\times10^{-2}$ | $[0.1305, 4.4671] \times 10^{-2}$ |
| $K_T$ | 38.6140 | $[4.3513, 93.0632]$ |
| $\lambda_D$ | 7.1822 | $[2.2653, 9.8756]$ |
| $K_I$ | 0.1379 | $[0.0240, 0.4024]$ |

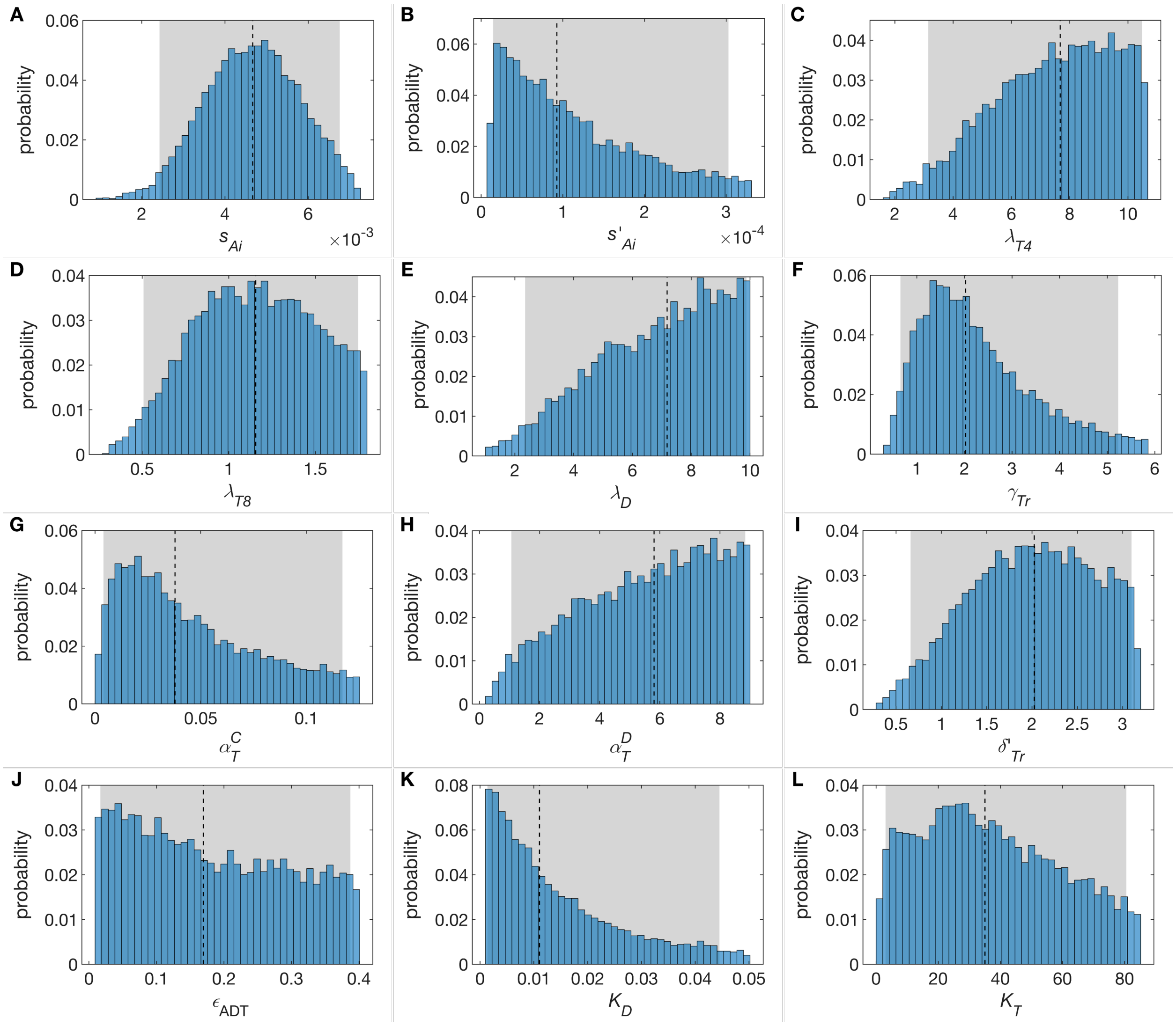


Supplementary Figure S2.1: Posterior distributions of parameters relating to immune presence in the tumor were inferred using a sample importance resampling algorithm. The resultant distributions, together with 95% confidence bounds (shaded gray areas) and median values (dashed black lines), are plotted.

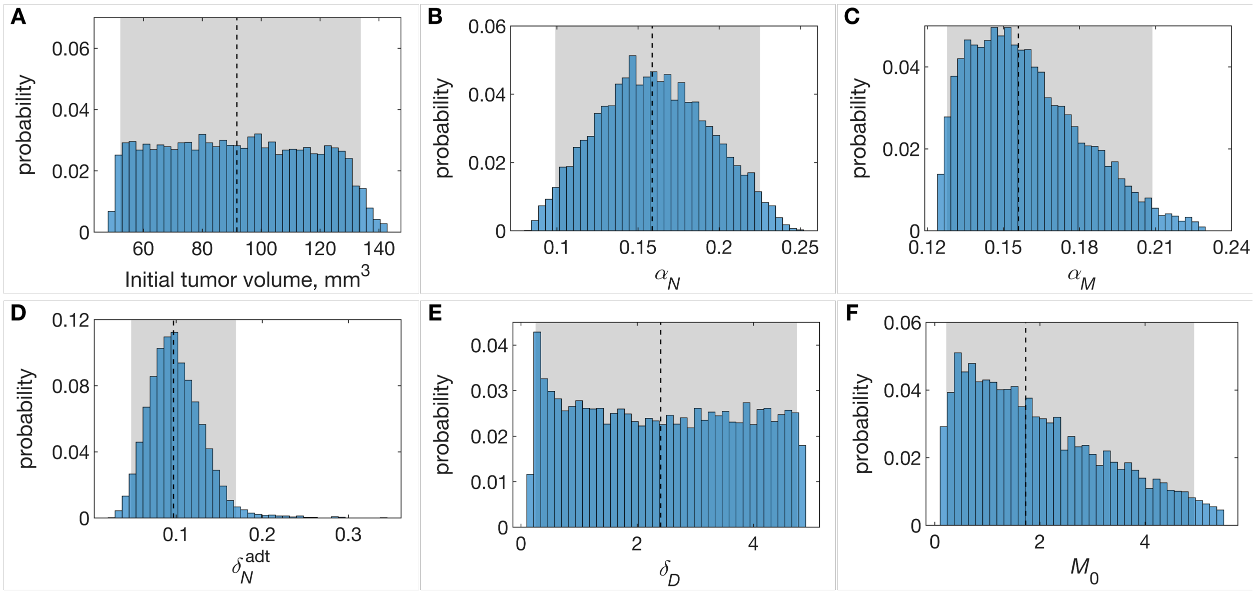


Supplementary Figure S2.2: Posterior distributions of parameters relating to tumor growth and response to ADT were inferred using a sample importance resampling algorithm. The resultant distributions, together with 95% confidence bounds (shaded gray areas) and median values (dashed black lines), are plotted.

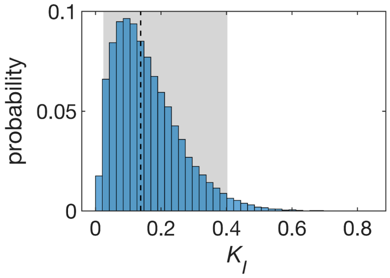


Supplementary Figure S2.3: Assumed distribution of $K_I$, the CTL to target cell ratio at which rate of CTL-induced cell kill is half its maximum value. Also shown are 95% confidence bounds (shaded gray area) and its median value (dashed black line).

#### S3 Approximation of Sensitivity Analysis

We identify those parameters that are significantly associated with a cure (tumor size $< 1$ mm$^3$), when mice are treated with combination ADT and vaccine per the optimal protocol suggested by our model. Table S3.1 gives trends in model parameters significantly associated with this outcome in an individual simulated mouse. Multiple logistic regression compared 318 cured mice to 4,559 'euthanized' or dead mice, with mice surviving-but-not-cured at the end of the simulation excluded from this analysis. Positive effect sizes indicate parameters that assume higher values in the cured mice, while negative effect sizes indicate the opposite. The p-values can be made arbitrarily small by increasing the number of simulated mice, but the relative value of Z-scores between parameters reflects their contribution to the outcome, approximating a sensitivity analysis if the model could be analyzed in such a way. Non-significant parameters are not reported in the table, for the sake of brevity.

Supplementary Table S3.1: **Model parameters significantly associated with cure under optimized combination therapy.**

| **Parameter** | **Dimensions** | **Effect Size** | **SE** | **Z** | **p-val** |
|---|---|---|---|---|---|
| $v_{tum}(10)$ | mm$^3$ | $-0.03595$ | 0.00559 | $-6.42890$ | $< 0.0001$ |
| $\alpha_N$ | 1/day | $-28.20485$ | 4.00701 | $-7.03888$ | $< 0.0001$ |
| $\alpha_M$ | 1/day | $-92.53038$ | 8.52813 | $-10.85002$ | $< 0.0001$ |
| $\delta_D$ | 1/day | 2.48784 | 0.18042 | 13.78921 | $< 0.0001$ |
| $M(10)$ | mm$^3$ | $-1.34657$ | 0.13234 | $-10.17497$ | $< 0.0001$ |
| $\lambda_{T8}$ | 1/day | 3.71596 | 0.42951 | 8.65152 | $< 0.0001$ |
| $\alpha_T^C$ | $\frac{\text{ng}}{10^6\text{cells}} \frac{1}{\text{day}}$ | $-47.28937$ | 5.16021 | $-9.16424$ | $< 0.0001$ |
| $\alpha_T^D$ | $\frac{\text{ng}}{10^6\text{cells}} \frac{1}{\text{day}}$ | $-1.85225$ | 0.12557 | $-14.75059$ | $< 0.0001$ |
| $\delta'_{Tr}$ | 1/day | 2.78649 | 0.26042 | 10.70020 | $< 0.0001$ |
| $K_T$ | $10^6$ cells/mm$^3$ | $-0.06876$ | 0.00715 | $-9.61972$ | $< 0.0001$ |
| $K_I$ | dimensionless | $-53.73040$ | 3.96685 | $-13.54485$ | $< 0.0001$ |

### References


[1] S. C. McKarns and R. H. Schwartz, "Distinct effects of tgf-$\beta$1 on cd4+ and cd8+ t cell survival, division, and il-2 production: a role for t cell intrinsic smad3," *The Journal of Immunology*, vol. 174, no. 4, pp. 2071–2083, 2005.

[2] M. Robertson-Tessi, A. El-Kareh, and A. Goriely, "A mathematical model of tumor–immune interactions," *Journal of theoretical biology*, vol. 294, pp. 56–73, 2012.

[3] A. Boissonnas, L. Fetler, I. S. Zeelenberg, S. Hugues, and S. Amigorena, "In vivo imaging of cytotoxic t cell infiltration and elimination of a solid tumor," *Journal of Experimental Medicine*, vol. 204, no. 2, pp. 345–356, 2007.

[4] K.-X. Zhang, J. Firus, B. Prieur, W. Jia, and P. S. Rennie, "To die or to survive, a fatal question for the destiny of prostate cancer cells after androgen deprivation therapy," *Cancers*, vol. 3, no. 2, pp. 1498–1512, 2011.

[5] M. R. Elliott and K. S. Ravichandran, "The dynamics of apoptotic cell clearance," *Developmental Cell*, vol. 38, no. 2, pp. 147–160, 2016.

[6] A. F. Maree, M. Komba, C. Dyck, M. Labecki, D. T. Finegood, and L. Edelstein-Keshet, "Quantifying macrophage defects in type 1 diabetes," *Journal of Theoretical Biology*, vol. 233, no. 4, pp. 533–551, 2005.

[7] B. Sauter, M. L. Albert, L. Francisco, M. Larsson, S. Somersan, and N. Bhardwaj, "Consequences of cell death: exposure to necrotic tumor cells, but not primary tissue cells or apoptotic cells, induces the maturation of immunostimulatory dendritic cells," *Journal of Experimental Medicine*, vol. 191, no. 3, pp. 423–434, 2000.

[8] A. Gardner and B. Ruffell, "Dendritic cells and cancer immunity," *Trends in Immunology*, vol. 37, no. 12, pp. 855–865, 2016.

[9] N. Larmonier, M. Marron, Y. Zeng, J. Cantrell, A. Romanoski, M. Sepassi, S. Thompson, X. Chen, S. Andreansky, and E. Katsanis, "Tumor-derived cd4+ cd25+ regulatory t cell suppression of dendritic cell function involves tgf-$\beta$ and il-10," *Cancer Immunology, Immunotherapy*, vol. 56, no. 1, pp. 48–59, 2007.

[10] J. Banchereau and R. M. Steinman, "Dendritic cells and the control of immunity," *Nature*, vol. 392, no. 6673, pp. 245–252, 1998.

[11] S. Halle, O. Halle, and R. Förster, "Mechanisms and dynamics of t cell-mediated cytotoxicity in vivo," *Trends in Immunology*, vol. 38, no. 6, pp. 432–443, 2017.

[12] S. Mitra and W. J. Leonard, "Biology of il-2 and its therapeutic modulation: mechanisms and strategies," *Journal of Leukocyte Biology*, vol. 103, no. 4, pp. 643–655, 2018.

[13] X. Ren, F. Ye, Z. Jiang, Y. Chu, S. Xiong, and Y. Wang, "Involvement of cellular death in trail/dr5-dependent suppression induced by cd4+ cd25+ regulatory t cells," *Cell Death and Differentiation*, vol. 14, no. 12, pp. 2076–2084, 2007.

[14] V. C. Liu, L. Y. Wong, T. Jang, A. H. Shah, I. Park, X. Yang, Q. Zhang, S. Lonning, B. A. Teicher, and C. Lee, "Tumor evasion of the immune system by converting cd4+ cd25- t cells into cd4+ cd25+ t regulatory cells: role of tumor-derived tgf-$\beta$," *The Journal of Immunology*, vol. 178, no. 5, pp. 2883–2892, 2007.

[15] Y. Y. Wan and R. A. Flavell, "Identifying foxp3-expressing suppressor t cells with a bicistronic reporter," *Proceedings of the National Academy of Sciences*, vol. 102, no. 14, pp. 5126–5131, 2005.

[16] Y. Y. Wan and R. A. Flavell, "'yin–yang' functions of transforming growth factor-$\beta$ and t regulatory cells in immune regulation," *Immunological Reviews*, vol. 220, no. 1, pp. 199–213, 2007.

[17] R. Alonso, H. Flament, S. Lemoine, C. Sedlik, E. Bottasso, I. Péguillet, V. Prémel, J. Denizeau, M. Salou, A. Darbois, *et al.*, "Induction of anergic or regulatory tumor-specific cd4+ t cells in the tumor-draining lymph node," *Nature Communications*, vol. 9, no. 1, pp. 1–7, 2018.

[18] M. Yadav, J. A. Bluestone, and S. Stephan, "Peripherally induced tregs–role in immune homeostasis and autoimmunity," *Frontiers in Immunology*, vol. 4, p. 232, 2013.

[19] E. Batlle and J. Massagué, "Transforming growth factor-$\beta$ signaling in immunity and cancer," *Immunity*, vol. 50, no. 4, pp. 924–940, 2019.

[20] B. Ludewig, P. Krebs, T. Junt, H. Metters, N. J. Ford, R. M. Anderson, and G. Bocharov, "Determining control parameters for dendritic cell-cytotoxic t lymphocyte interaction," *European Journal of Immunology*, vol. 34, no. 9, pp. 2407–2418, 2004.

[21] J. B. Wing, A. Tanaka, and S. Sakaguchi, "Human foxp3+ regulatory t cell heterogeneity and function in autoimmunity and cancer," *Immunity*, vol. 50, no. 2, pp. 302–316, 2019.

[22] M. Miyara, Y. Yoshioka, A. Kitoh, T. Shima, K. Wing, A. Niwa, C. Parizot, C. Taflin, T. Heike, D. Valeyre, *et al.*, "Functional delineation and differentiation dynamics of human cd4+ t cells expressing the foxp3 transcription factor," *Immunity*, vol. 30, no. 6, pp. 899–911, 2009.

[23] S. Yamazaki, T. Iyoda, K. Tarbell, K. Olson, K. Velinzon, K. Inaba, and R. M. Steinman, "Direct expansion of functional cd25+ cd4+ regulatory t cells by antigen-processing dendritic cells," *Journal of Experimental Medicine*, vol. 198, no. 2, pp. 235–247, 2003.

[24] C.-L. Tran, A. D. Jones, and K. Donaldson, "Mathematical model of phagocytosis and inflammation after the inhalation of quartz at different concentrations.," *Scandinavian Journal of Work, Environment & Health*, vol. 21, pp. 50–54, 1995.

[25] J. A. Borghans, R. J. De Boer, and L. A. Segel, "Extending the quasi-steady state approximation by changing variables," *Bulletin of Mathematical Biology*, vol. 58, no. 1, pp. 43–63, 1996.

[26] D. Coombs, O. Dushek, and P. A. van der Merwe, "A review of mathematical models for t cell receptor triggering and antigen discrimination," in *Mathematical Models and Immune Cell Biology* (C. Molina-París and G. Lythe, eds.), pp. 25–45, New York, NY: Springer, 2011.

[27] R. J. De Boer and A. S. Perelson, "Towards a general function describing t cell proliferation," *Journal of Theoretical Biology*, vol. 175, no. 4, pp. 567–576, 1995.

[28] S. Schnell, "Validity of the michaelis–menten equation–steady-state or reactant stationary assumption: that is the question," *The FEBS journal*, vol. 281, no. 2, pp. 464–472, 2014.

[29] S. Park, R. R. Ang, S. P. Duffy, J. Bazov, K. N. Chi, P. C. Black, and H. Ma, "Morphological differences between circulating tumor cells from prostate cancer patients and cultured prostate cancer cells," *PloS one*, vol. 9, no. 1, p. e85264, 2014.

[30] E. Maeno, T. Tsubata, and Y. Okada, "Apoptotic volume decrease (avd) is independent of mitochondrial dysfunction and initiator caspase activation," *Cells*, vol. 1, no. 4, pp. 1156–1167, 2012.

[31] Y. Okada, E. Maeno, T. Shimizu, K. Manabe, S.-i. Mori, and T. Nabekura, "Dual roles of plasmalemmal chloride channels in induction of cell death," *Pflügers Archiv*, vol. 448, no. 3, pp. 287–295, 2004.

[32] H. Pasantes-Morales, "Channels and volume changes in the life and death of the cell," *Molecular pharmacology*, vol. 90, no. 3, pp. 358–370, 2016.

[33] M. J. Miller, A. S. Hejazi, S. H. Wei, M. D. Cahalan, and I. Parker, "T cell repertoire scanning is promoted by dynamic dendritic cell behavior and random t cell motility in the lymph node," *Proceedings of the National Academy of Sciences*, vol. 101, no. 4, pp. 998–1003, 2004.

[34] E. H. Chapman, A. S. Kurec, and F. Davey, "Cell volumes of normal and malignant mononuclear cells.," *Journal of Clinical Pathology*, vol. 34, no. 10, pp. 1083–1090, 1981.

[35] R. Kushwah and J. Hu, "Dendritic cell apoptosis: regulation of tolerance versus immunity," *The Journal of Immunology*, vol. 185, no. 2, pp. 795–802, 2010.

[36] A. Lanzavecchia and F. Sallusto, "Regulation of t cell immunity by dendritic cells," *Cell*, vol. 106, no. 3, pp. 263–266, 2001.

[37] M. Berard and D. F. Tough, "Qualitative differences between naive and memory t cells," *Immunology*, vol. 106, no. 2, pp. 127–138, 2002.

[38] A. Radunskaya, R. Kim, and T. Woods II, "Mathematical modeling of tumor immune interactions: a closer look at the role of a pd-l1 inhibitor in cancer immunotherapy," *Spora: A Journal of Biomathematics*, vol. 4, no. 1, pp. 25–41, 2018.

[39] C. T. Luo, W. Liao, S. Dadi, A. Toure, and M. O. Li, "Graded foxo1 activity in t reg cells differentiates tumour immunity from spontaneous autoimmunity," *Nature*, vol. 529–536, no. 7587, p. 532, 2016.

[40] B. Breart, F. Lemaître, S. Celli, and P. Bousso, "Two-photon imaging of intratumoral cd8+ t cell cytotoxic activity during adoptive t cell therapy in mice," *The Journal of Clinical Investigation*, vol. 118, no. 4, pp. 1390–1397, 2008.

[41] L. M. Wakefield, T. Winokur, R. Hollands, K. Christopherson, A. Levinson, and M. Sporn, "Recombinant latent transforming growth factor beta 1 has a longer plasma half-life in rats than active transforming growth factor beta 1, and a different tissue distribution.," *The Journal of Clinical Investigation*, vol. 86, no. 6, pp. 1976–1984, 1990.

[42] A. Alloatti, F. Kotsias, J. G. Magalhaes, and S. Amigorena, "Dendritic cell maturation and cross-presentation: timing matters!," *Immunological Reviews*, vol. 272, no. 1, pp. 97–108, 2016.

[43] G. J. Randolph, V. Angeli, and M. A. Swartz, "Dendritic-cell trafficking to lymph nodes through lymphatic vessels," *Nature Reviews Immunology*, vol. 5, no. 8, pp. 617–628, 2005.

[44] Y.-C. Shen, A. Ghasemzadeh, C. M. Kochel, T. R. Nirschl, B. J. Francica, Z. A. Lopez-Bujanda, M. A. C. Haro, A. Tam, R. A. Anders, M. J. Selby, A. J. Korman, and C. G. Drake, "Combining intratumoral treg depletion with androgen deprivation therapy (adt): preclinical activity in the myc-cap model," *Prostate Cancer and Prostatic Diseases*, vol. 21, no. 1, pp. 113–125, 2018.

[45] S. Wei, I. Kryczek, and W. Zou, "Regulatory t-cell compartmentalization and trafficking," *Blood*, vol. 108, no. 2, pp. 426–431, 2006.

[46] R. J. De Boer, D. Homann, and A. S. Perelson, "Different dynamics of cd4+ and cd8+ t cell responses during and after acute lymphocytic choriomeningitis virus infection," *The Journal of Immunology*, vol. 171, no. 8, pp. 3928–3935, 2003.

[47] T. R. Mempel, S. E. Henrickson, and U. H. Von Andrian, "T-cell priming by dendritic cells in lymph nodes occurs in three distinct phases," *Nature*, vol. 427, no. 6970, p. 154, 2004.

[48] B. J. Rollins, T. M. O'Connell, G. Bennett, L. E. Burton, C. D. Stiles, and J. G. Rheinwald, "Environment-dependent growth inhibition of human epidermal keratinocytes by recombinant human transforming growth factor-beta," *Journal of Cellular Physiology*, vol. 139, no. 3, pp. 455–462, 1989.

[49] L. Ellis, K. Lehet, S. Ramakrishnan, R. Adelaiye, and R. Pili, "Development of a castrate resistant transplant tumor model of prostate cancer," *The Prostate*, vol. 72, no. 6, pp. 587–591, 2012.

[50] F. Bladou, R. L. Vessella, K. R. Buhler, W. J. Ellis, L. D. True, and P. H. Lange, "Cell proliferation and apoptosis during prostatic tumor xenograft involution and regrowth after castration," *International Journal of Cancer*, vol. 67, no. 6, pp. 785–790, 1996.

[51] C. K. Søgaard, S. A. Moestue, M. B. Rye, J. Kim, A. Nepal, N.-B. Liabakk, S. Bachke, T. F. Bathen, M. Otterlei, and D. K. Hill, "Apim-peptide targeting pcna improves the efficacy of docetaxel treatment in the tramp mouse model of prostate cancer," *Oncotarget*, vol. 9, no. 14, pp. 11752–11766, 2018.

[52] F. Famoye, E. Akarawak, and M. Ekum, "Weibull-normal distribution and its applications," *Journal of Statistical Theory and Applications*, vol. 17, no. 4, pp. 719–727, 2018.
